# AbPACER: parent-aware, affinity-label-blind prioritization of affinity-matured scFv clones from phage-display NGS

**DOI:** 10.64898/2026.08.05.742956

**Authors:** Ahn Jae Chung, Byung Young Park, Eun-Byeol Park, Jae-Ho Han

## Abstract

**Background:** Affinity-maturation phage-display next-generation sequencing (NGS) yields more paired single-chain variable fragment clones than can be characterized experimentally, creating a fixed-budget prioritization problem. Read counts provide empirical support rather than direct affinity labels. We developed AbPACER (Antibody Parent-Aware Contextual Evidence Ranker), an affinity-label-blind neural ranker combining parent-relative mutation descriptors, frozen antibody-language-model context, and NGS evidence from related clones. AbPACER is campaign-adaptive rather than zero-shot: for each campaign, it is fitted to paired sequences and round-resolved R1–R3 counts before returning a 384-candidate assay list. We evaluated it in two retrospective phage-display campaigns and separately assessed its supervised mean-squared-error adaptation on AlphaSeq, denoted AbPACER-MSE.

**Results:** From frozen top-5% candidate sets containing 16,323 Fas-associated factor 1 (FAF1) and 7,487 vascular endothelial growth factor receptor (VEGFR) clones, each method ranked the complete target-specific set and selected 384 candidates. In FAF1, AbPACER recovered 2.00 ± 0.00 of seven retrospective panel clones, recovering two in every seed, compared with 1*/*7 by total count, 1.00 ± 0.00 by Ens-Grad CNN, 1.67 ± 1.15 by A2Binder-HL, and 1.33 ± 0.58 by AbAffinity. In VEGFR, AbPACER recovered 2.33 ± 0.58 of three panel clones, the highest observed learned-method mean, whereas total count recovered 3*/*3. No learned method was uniformly best at broader hypothetical budgets. On the public AlphaSeq common split of 11,670 fixed-test variants, AbPACER-MSE recovered 187.0 ± 2.6 of the true top-384, closely matching AbAffinity (188.0 ± 2.6) and exceeding A2Binder (175.7 ± 6.4) and Ens-Grad CNN (154.0 ± 6.1). AbPACER-MSE updated 1.378 million task-specific parameters, compared with 651.04 million for AbAffinity, and achieved Pearson 0.687 ± 0.003 and Spearman 0.652 ± 0.002.

**Conclusions:** AbPACER provides a campaign-specific, parent-aware framework for fixed-budget prioritization from affinity-label-blind phage-display NGS data. At the 384-candidate endpoint, it showed the highest mean recovery among learned methods in both retrospective campaigns. AbPACER-MSE closely matched AbAffinity in true top-384 recovery while updating substantially fewer task-specific parameters. These results motivate prospective evaluation of sequence-conditioned reranking as a complement to count-based prioritization.

## 1 Background

Antibody phage display links genotype to binding phenotype and is widely used for in vitro antibody discovery and affinity maturation [1, 2, 3]. Iterative selection generates large families of related variants, while next-generation sequencing (NGS) reveals these pools at substantially greater depth than conventional colony picking [4, 5]. Long-read sequencing can additionally recover full-length single-chain variable fragment (scFv) repertoires while preserving the linked variable heavy (VH) and variable light (VL) regions within each construct [6]. Hundreds of thousands of paired VH/VL clones may therefore be observed, although expression, purification, and quantitative binding characterization remain feasible for only a small fraction. The practical computational task is consequently not sequence generation alone, but prioritization of observed candidates under a fixed assay budget. The required output is an auditable assay-prioritization list rather than an unconstrained sequence score.

Total read count is a strong operational signal because it summarizes cumulative empirical support from observed clone abundance across panning outputs. It is not, however, a direct measure of molecular affinity and may also reflect clone-dependent phage amplification [7], as well as variation in host fitness, protein expression, display, folding, sequencing, and sampling [8]. In this study, total count was therefore treated as affinity-label-blind empirical evidence rather than as a measured affinity label: it defined the count-supported candidate universe, served as the primary deterministic baseline, and contributed to the NGS-derived supervisory target used for model fitting. A count-based filter can remove the lowest-support sequencing tail, but the resulting candidate universe may still contain thousands of clones and large exact-count tie groups. The remaining computational question is therefore whether parent-aware sequence modeling can extract a more useful ordering from cumulative count and panning-trajectory evidence than direct ranking by those scalar signals alone.

Previous computational studies have addressed related but distinct tasks. Enrichment-trained models have guided complementarity-determining region (CDR) optimization [9], and generative models have learned enriched repertoire structure [10]. Supervised models have predicted quantitative affinity or related binding properties from experimentally labeled antibody libraries [11]. AlphaSeq provides a densely measured public landscape for model development and benchmarking [12], on which the recent AbAffinity preprint evaluated language-model-based affinity prediction [13]. Recent Bayesian preprint work has also modeled experimental noise and uncertainty in phage-display data [14]. These studies motivate sequence-aware analysis but do not directly address affinity-label-blind prioritization of already observed, paired VH/VL affinity-matured clones under a fixed experimental budget. This setting motivates modeling candidates relative to their parental clone, deriving ranking supervision without affinity labels, and directly ordering all candidates in a frozen assay universe without treating unmeasured clones as affinity negatives.

Affinity maturation provides a natural reference because each candidate is a mutational departure from a known parental clone and the effect of a mutation depends on its sequence background [15, 16]. AbPACER preserves paired VH/VL identity and combines explicit parent-to-candidate mutation descriptors with frozen IgBERT residue-level and global sequence context [17]. It learns a global pairwise ordering from an affinity-label-blind target combining cumulative count and panning trajectory, aggregates NGS evidence from related unmeasured clones, and applies a bounded local-family correction. The resulting AbPACER score directly ranks candidates for experimental selection rather than estimating a calibrated affinity value.

AbPACER is intended as a campaign-adaptive prioritizer rather than a zero-shot predictor transferred unchanged across campaigns. For each new affinity-maturation campaign, the model is fitted de novo using the available pool of paired VH/VL sequences and round-resolved R1–R3 counts before quantitative affinity characterization. It then ranks the frozen count-supported candidate manifest and returns the top 384 candidates for experimental assay. The objective is therefore to allocate limited screening capacity using all affinity-label-blind information available within the current campaign.

The study contains two complementary evaluation arms organized around fixed-budget prioritization. The retrospective phage-display arm evaluates affinity-label-blind ranking in two affinity-maturation campaigns, whereas the public AlphaSeq landscape evaluates supervised adaptation of the same parent-aware representation under dense affinity labels [12]. Across both arms, the primary selection depth was fixed before recovery evaluation at 384 candidates as an operational high-throughput downstream-screening benchmark motivated by established 384-well antibody-screening and immunoassay formats [18, 19]. This candidate-level budget was distinct from the pooled panning format used to generate the R1–R3 counts. Applying the same selection depth to AlphaSeq provided a common operational endpoint rather than a pooled biological benchmark; the two arms use distinct supervision, datasets, and recovery denominators, and their results are not pooled.

Here, we evaluate AbPACER in two retrospective affinity-maturation campaigns and separately assess the same parent-aware representation on AlphaSeq. The accompanying release will provide paired-chain data, complete rankings, and versioned reproducibility artifacts.

## 2 Materials and methods

### 2.1 Study design and dataset generation

The retrospective phage-display arm evaluated AbPACER and common-protocol comparators within frozen count-supported candidate sets. NGS-matched clone keys with quantitative measurements were excluded from split construction, fitting, internal validation, neighbor-reference pools, and checkpoint selection, but their paired VH/VL sequences and R1–R3 counts remained in the candidate pools for scoring. ELISA-derived apparent-affinity values were used only for retrospective evaluation.

For deployment, AbPACER is fitted separately for each campaign using eligible paired VH/VL sequences and round-resolved R1–R3 counts available before quantitative affinity characterization, and then ranks the frozen count-supported manifest to produce a 384-candidate assay list. In the retrospective analyses, quantitatively measured clones served as evaluation outcomes while remaining excluded from all fitting and model-selection steps.

AbPACER-MSE separately evaluated the same parent-aware representation on AlphaSeq [12]. It used neither phage-display counts or trajectories nor the phage-specific target or neighbor evidence; the two arms used distinct supervision and were analyzed separately.

AngioLab, Inc. generated the Fas-associated factor 1 (FAF1) and vascular endothelial growth factor receptor (VEGFR) affinity-maturation datasets in separate campaigns using a human naive scFv phage-display library. FAF1_#1 and VEGFR clone #163 were selected as the respective parental clones for random-mutagenesis affinity maturation. The VEGFR lineage was also characterized against VEGFR-1 for descriptive purposes, whereas the panning outputs and NGS counts analyzed here were generated by selection against VEGFR-2. Parent-derived libraries underwent three rounds of increasingly stringent solid-phase panning, and PacBio Revio long-read sequencing of full-length scFv amplicons recovered linked VH and VL sequences from the same construct. Phage-display prioritization used the paired VH/VL sequences and R1–R3 panning-output counts; additional experimental details are provided in Additional file 1.

In a separate experimental branch, R3 colonies were randomly picked and screened by single-chain variable fragment–Fc (scFv–Fc) binding ELISA. Screen-advanced candidates were expressed as scFv– Fc proteins and evaluated by concentration-dependent binding ELISA. Only screen-advanced clones underwent quantitative measurement. All quantitative ELISA results generated in this workflow were included in the reported set, and no quantitatively measured clone was excluded on the basis of its binding result. The fitted midpoint was reported as an ELISA-derived apparent-affinity value, with lower values indicating stronger apparent binding. Available curve-level fit outputs are provided in Additional file 1, Fig. S1. Because the campaigns and measured panels were retrospective, the panels were not exhaustive catalogs of favorable variants, and neither campaign constituted an independent external test. Evaluation therefore quantified recovery of evaluation-panel clones at fixed assay budgets rather than full-pool sensitivity or specificity. For the phage comparisons, all learned methods used the same frozen candidate manifests and fixed train/validation memberships across seeds 42–44. Across-seed summaries reflected optimization variability rather than biological replication, and no retrospective recovery result was used to reselect a seed, training duration, split, or candidate universe.

The experimental workflow and the separation of the NGS-ranking and retrospective ELISA-evaluation branches are summarized in Fig. 1.

**Figure 1:**
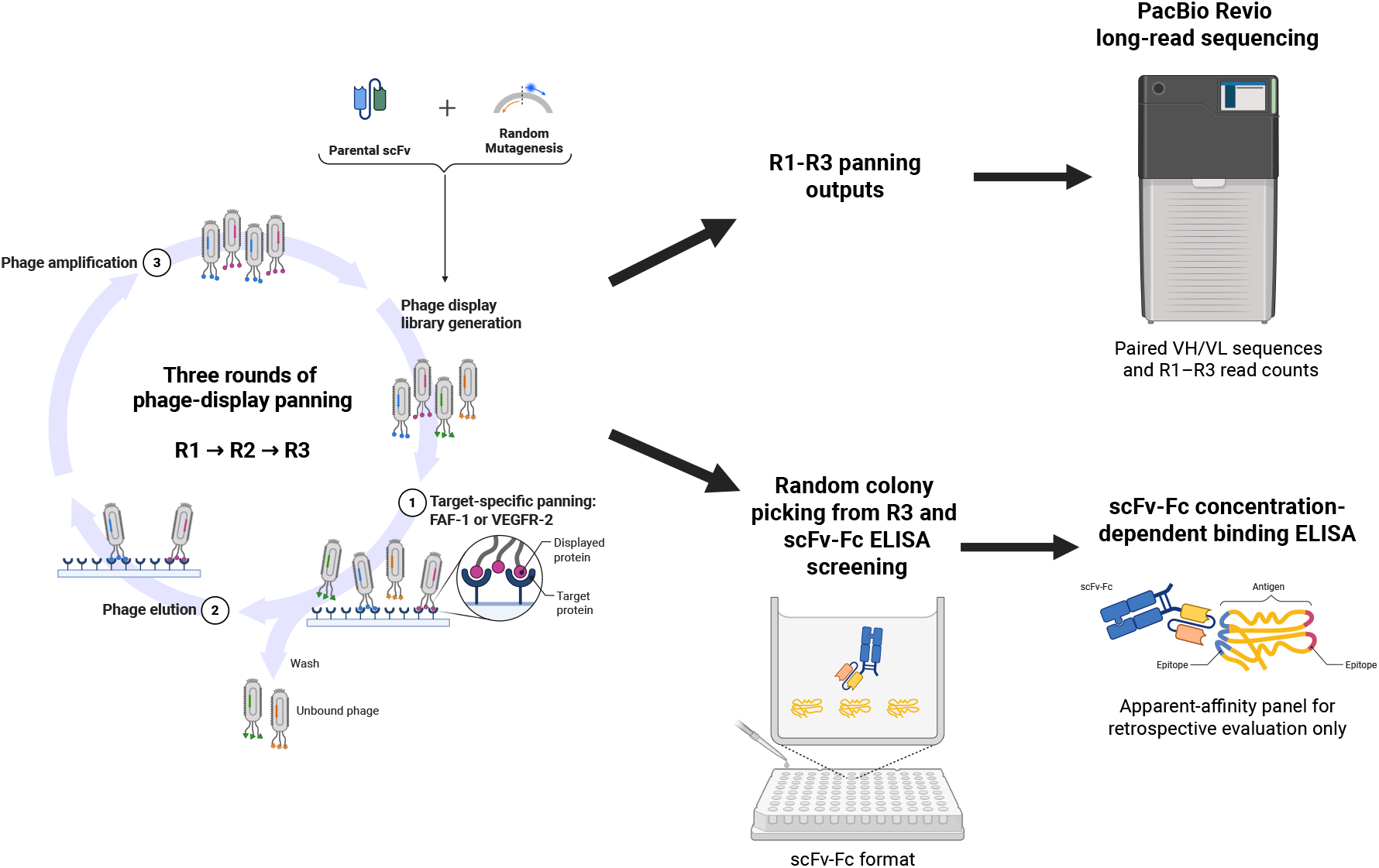
Experimental generation and use of the affinity-maturation phage-display datasets. Parental scFv sequences were diversified and subjected to three rounds of target-specific panning. PacBio Revio sequencing recovered paired VH/VL sequences and R1–R3 read counts for affinity-label-blind fitting and ranking. In a separate experimental branch, R3 colonies were screened by scFv–Fc binding ELISA, and screen-advanced candidates were expressed as scFv–Fc proteins for concentration-dependent binding ELISA. The resulting ELISA-derived apparent affinity measurements formed the available quantitative measurement set; the fixed retrospective evaluation panels were then defined by exact NGS-key matching and membership in the frozen candidate set. These values did not enter fitting, checkpoint selection, score computation, or within-method candidate ordering; the corresponding paired sequences and R1–R3 counts remained in the candidate sets and were ranked with all other candidates. Created with BioRender.com [20].

**Figure 2:**
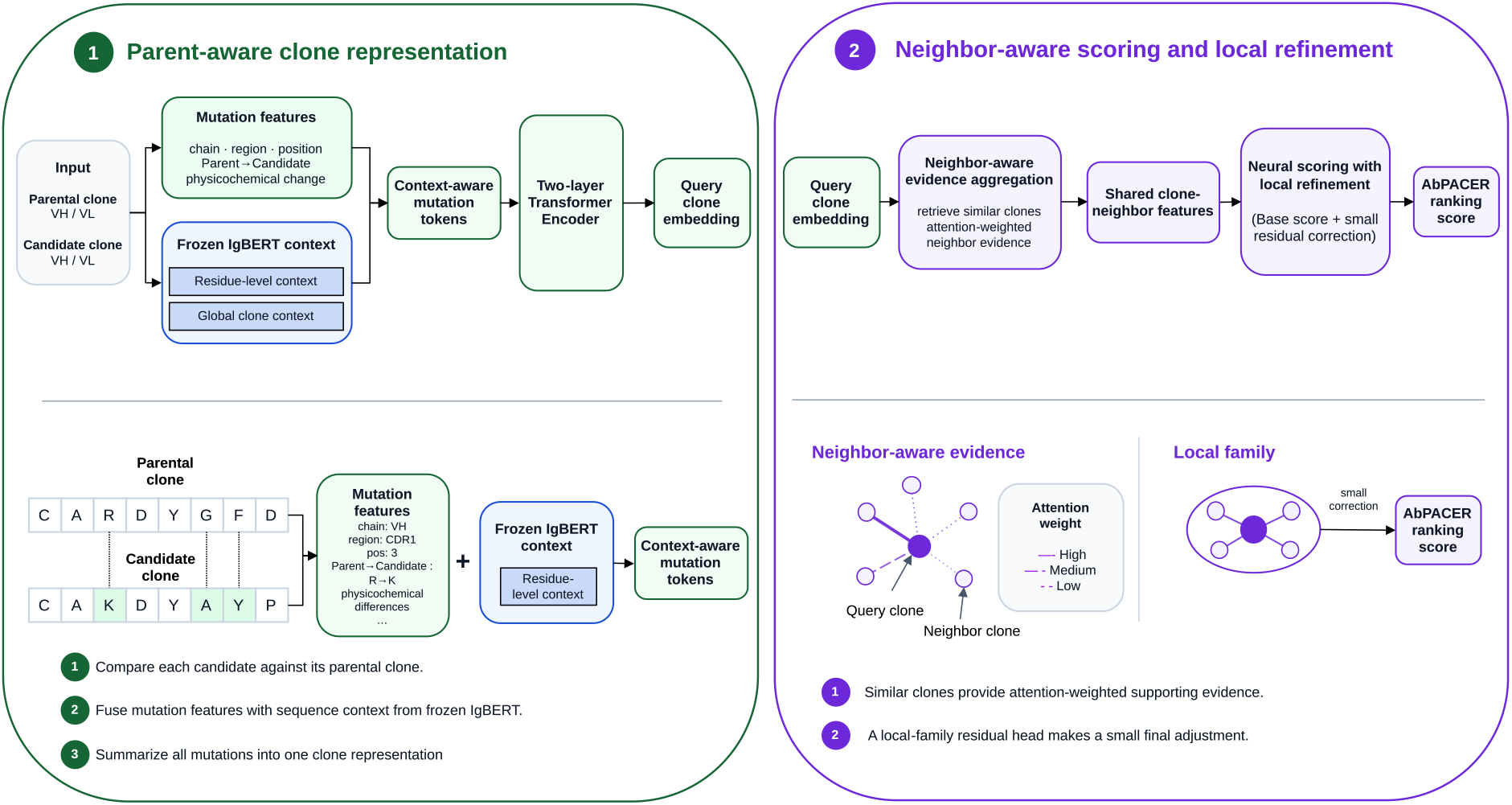
Overview of AbPACER. Each candidate is represented as a paired VH/VL change from its target-specific parental clone using explicit mutation descriptors and frozen IgBERT residue-level and global sequence context. A transformer integrates the variable-length mutation set into the fixed-length query representation *q_i_*. Unmeasured reference clones retrieved in parent-relative IgBERT-difference space contribute attention-weighted NGS evidence to the neighbor-aware base score *b_i_*. A separate local-family graph supplies the ordering signal for the bounded residual correction *δ_i_*, yielding the final AbPACER ranking score *s_i_* = *b_i_* + *δ_i_*. Candidates are ranked directly by decreasing *s_i_* for fixed-budget selection.

### 2.2 Preprocessing, candidate sets, and retrospective evaluation

PacBio reads were translated from the detected start methionine, and paired VH and VL segments were parsed using the known scFv linker. Reads with frameshifts, incomplete translations, unparseable paired chains, or premature stop codons outside the designed amber-suppressible context were removed. Designed NNK-derived TAG codons in the amber-suppressible context were translated as glutamine (Q). Retained reads were collapsed by exact paired amino-acid key, VH|VL, and duplicate-key counts were summed separately for R1, R2, and R3. Total read count was defined as

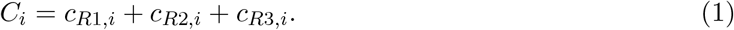

Total count was treated as cumulative affinity-label-blind empirical read support rather than as molecular affinity. It defined the candidate sets, served as the primary non-learned baseline, and contributed, together with panning-trajectory evidence, to the affinity-label-blind fitting target defined in the following subsection. Candidate universes were constructed by total count and frozen before comparative evaluation, with all clone keys tied at a cutoff retained and the exact memberships recorded as reproducibility manifests.

The top-5% candidate manifests were designated as the primary operational setting independently of ELISA-derived apparent-affinity values or recovery outcomes. They removed the extensive low-count sequencing tail while retaining 16,323 FAF1 and 7,487 VEGFR candidates, with realized minimum total counts of 6 and 18, respectively. These sets remained approximately 42.5-fold and 19.5-fold larger than the 384-candidate selection depth, leaving a substantial reranking problem. The top-1% and top-10% manifests are reported descriptively in Additional file 1 to document neighboring candidate-universe scales and panel coverage. Full-pool scale, singleton burden, primary-candidate-set size, and retrospective evaluation-panel construction are summarized in Table 1.

**Table 1:** Full-pool phage-display statistics, frozen primary candidate sets, and retrospective evaluation design.

| Target | Unique paired VH/VL<br>clone keys | R1<br>reads | R2<br>reads | R3<br>reads | Total<br>reads | Total count = 1,<br>n (%) |
| --- | --- | --- | --- | --- | --- | --- |
| FAF1 | 327 258 | 128 445 | 472 725 | 117 998 | 719 168 | 206 584 (63.1%) |
| VEGFR-2 | 148 073 | 216 181 | 489 642 | 449 405 | 1 155 228 | 79 708 (53.8%) |

**Panel B. Frozen primary candidate sets and retrospective evaluation panels**
| Target | Parental clone and<br>apparent affinity | Top-5%<br>candidate<br>set | Quantitatively measured /<br>exact NGS match /<br>primary panel | Primary-panel<br>apparent-affinity<br>range |
| --- | --- | --- | --- | --- |
| FAF1 | FAF1_#1 (43.1 pM) | 16 323 | 9/8/7 | 21.8–35.3 pM |
| VEGFR-2 | #163 (20.7 nM) | 7 487 | 3/3/3 | 1.77–6.54 nM |

Panel A summarizes the complete R1–R3 sequencing pools after preprocessing and collapsing by exact paired VH/VL amino-acid key. “Total count = 1” means that a paired clone key had a cumulative R1–R3 total read count of one. Panel B reports the top-5% candidate sets, which were frozen before comparative evaluation; all clones tied at a cutoff count were retained. Quantitatively measured/exact NGS match/primary panel gives the numbers of clones with available quantitative scFv–Fc concentration–response ELISA measurements, exact collapsed paired VH/VL matches, and membership in the fixed primary evaluation panel, respectively. ELISA-derived apparent-affinity values were fitted midpoints from concentration-dependent scFv–Fc binding ELISA against the selection antigen, with lower values indicating stronger apparent binding. VEGFR-1 measurements were descriptive and did not define target-specific recovery.

**Table 2:** Primary evaluation-panel recovery after selecting 384 candidates from each frozen target-specific manifest.

| Selection rule or method | Task-specific trainable parameters | FAF1<br>384 of 16 323 selected<br>panel recovery (of 7) | VEGFR<br>384 of 7 487 selected<br>panel recovery (of 3) |
| --- | --- | --- | --- |
| Uniform random selection (expected) | – | 0.165 | 0.154 |
| Total read count | – | 1 | 3 |
| Peak trajectory | – | 1 | 3 |
| Direct $O_i$ ranking | – | 1 | 3 |
| Ens-Grad CNN | 0.058 M | $1.00 \pm 0.00$ | $0.33 \pm 0.58$ |
| A2Binder-HL | 114.63 M | $1.67 \pm 1.15$ | $1.00 \pm 0.00$ |
| AbAffinity | 651.04 M | $1.33 \pm 0.58$ | $1.33 \pm 0.58$ |
| AbPACER | 1.38 M | **$2.00 \pm 0.00$** | **$2.33 \pm 0.58$** |

Uniform-random values are the expected numbers of fixed-panel clones recovered when 384 candidates are sampled without replacement from each frozen manifest, $E[X] = 384K/N$ , where $K$ is the number of panel clones and $N$ is the manifest size. Learned values are mean ± sample standard deviation over seeds 42–44; deterministic controls have no stochastic variation. All methods ranked the complete frozen target-specific manifest using direct model outputs and the same deterministic score-tie rule. Parameter counts include task-specific trainable parameters and exclude frozen or non-trainable components. Boldface denotes the highest observed learned-method mean.

The Base stage used all eligible unmeasured rows from the campaign-wide collapsed pool, whereas the residual stage and final assay ranking were restricted to the frozen top-5% count-supported region. Thus, the top-5% manifest defined the operational assay universe rather than the complete source of fitting evidence.

Quantitative scFv–Fc concentration–response ELISA measurements were available for nine FAF1 clones and three VEGFR clones. One FAF1 clone did not match any exact collapsed paired VH|VL NGS key after final preprocessing. Of the remaining eight NGS-matched FAF1 clones, one had a total count of 4 and therefore fell outside the frozen primary top-5% candidate set. The remaining seven FAF1 clones constituted the fixed primary evaluation panel. All three VEGFR clones had exact collapsed paired VH|VL matches and were present in the primary candidate set. Among the quantitatively measured clones, panel inclusion was determined by exact NGS-key matching and membership in the frozen candidate set rather than by the measured apparent-affinity values. VEGFR-2 ELISA-derived apparent-affinity values defined target-specific recovery, whereas VEGFR-1 measurements were descriptive. Exact memberships, mutations, counts, and ranks are reported in Additional file 1.

The primary phage-display selection depth was fixed before evaluation-panel recovery at 384 candidate selections. This depth was chosen as a nominal high-throughput downstream-screening budget motivated by established 384-well antibody-screening and immunoassay formats [18, 19], rather than by the historical number of colonies screened. The 96-well format used during affinity-maturation panning did not define a 96-candidate assay budget because each panning well processed a pooled phage library rather than one individually assigned candidate clone. The value 384 therefore denotes a candidate-level computational assay budget rather than the well count of the upstream panning experiment or the exact occupancy of a single physical downstream plate; laboratory implementation may require additional wells or plates for controls and replicates. Reference and control entries were not included in candidate recovery. Budgets of 1,000, 2,000, and 5,000 represented secondary hypothetical scaling analyses within the primary manifest. An evaluation-panel clone was counted as recovered when its method-specific rank did not exceed the applicable budget. Because unmeasured candidates had unknown affinity and were not treated as negatives, recovery was a panel-based endpoint rather than an estimate of full-universe precision, recall, sensitivity, or specificity.

For each learned phage-display method, every candidate in the applicable frozen manifest was ranked directly by decreasing model score, with exact ties resolved by the frozen paired VH|VL key. NGS-matched quantitatively measured clone keys were excluded before split construction and did not enter fitting, internal validation, neighbor-reference pools, or checkpoint selection. Their paired VH/VL sequences and R1–R3 counts remained in the complete collapsed pools, and any such clone present in the applicable frozen candidate manifest was scored and ranked with all other candidates. No apparent-affinity value entered model fitting, checkpoint selection, score computation, or within-method candidate ordering.

To prevent closely related mutation patterns from crossing the fit and validation partitions, unmeasured candidates with identical parent-relative, chain-labeled mutation-position signatures were assigned to the same fixed partition. These memberships remained fixed across methods and seeds, and checkpoint selection used only affinity-label-blind targets from unmeasured validation rows. Exact split construction, software implementation, split sizes, and audit details are provided in Additional file 1.

A secondary frozen-output audit reused the six seed-specific final AbPACER direct-score rankings and the frozen top-5% manifests without new training, inference, checkpoint selection, calibration, post-ranking fusion, or consensus ranking. Exact-count tie intervals were calculated under descending total-count precedence. For the selection-pattern analysis, AbPACER-only candidates were those in the AbPACER top 384 but not the deterministic total-count top 384, whereas displaced candidates were the converse. Evaluation-panel clones were excluded from feature discovery, statistical testing, support filtering, and feature-stability assessment and were annotated only after the feature set had been frozen; complete definitions are provided in Additional file 1.

### 2.3 Affinity-label-blind NGS evidence

All learned phage-display methods used a common affinity-label-blind fitting target that combined cumulative empirical read support with relative abundance change across panning rounds. Neither component was an ELISA-derived apparent-affinity value, and their combination was treated as NGS evidence rather than a calibrated affinity estimate.

For clone *i* in round *r*, let *c_r,i_* be its read count, *C_r_* = ∑*_i_c_r,i_* the total read count in that round, and *N* the number of exact paired VH|VL keys in the collapsed R1–R3 union. A clone absent from a round was assigned *c_r,i_* = 0 before application of the fixed pseudocount *p* = 1. The smoothed relative frequency was

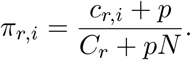

Normalization over the fixed clone universe adjusted for round-level sequencing-depth differences, so log ratios reflected changes in relative abundance rather than total read output. The pseudocount stabilized zero observations but did not correct clone-specific amplification, display, or sequencing biases.

Using natural logarithms, support-normalized enrichment from earlier round *a* to later round *b* was

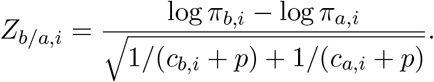

The denominator downweighted relative-abundance changes supported by few reads. *Z_b/a,i_* was therefore used as an operational support-normalized enrichment score rather than a calibrated inferential *z*-statistic; more positive values indicated stronger relative enrichment.

Peak trajectory was defined as

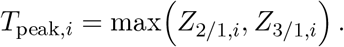

Round-to-round enrichment has precedent as an affinity-label-blind supervisory signal; Ens-Grad, for example, used the R2-to-R3 log-frequency ratio because that transition had the highest signal-to-noise ratio in its campaign [9]. Here, R1 was the common reference because it was the earliest sequenced panning output. Taking the maximum captured either enrichment already evident by R2 or enrichment that became strongest by R3, without requiring monotonic growth or assuming that one transition was uniformly most informative across FAF1 and VEGFR. *Z*_3_*_/_*_2_*_,i_*, representing only incremental change after R2, was retained as a separate deterministic baseline. Thus, *T*_peak_*_,i_* was an operational summary of R1-anchored selection-trajectory evidence rather than a kinetic model.

Before combination, total count was transformed as log(1 + *C_i_*) to limit the influence of its heavy upper tail while preserving its ordering. During holdout fitting, log(1 + *C_i_*) and *T*_peak_*_,i_* were standardized using target-specific means and standard deviations estimated from the fixed unmeasured training rows and applied unchanged to validation rows. For the final refit and candidate ranking, these moments were re-estimated from all eligible unmeasured rows and applied to the frozen candidate manifest. The global NGS-derived target was

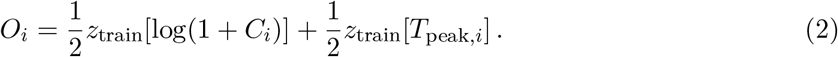

Cumulative count and peak trajectory captured complementary aspects of the panning observations: the former represented empirical support accumulated across R1–R3, whereas the latter represented directional change in relative abundance from R1. After standardization placed them on comparable numerical units, the equal coefficients imposed a fixed symmetric 1:1 weighting, so a one-training-set-standard-deviation increase in either component contributed 0.5 to *O_i_*. The factor 1*/*2 therefore made *O_i_*the arithmetic mean of the two standardized evidence coordinates. Because no measured affinity labels were available within the fitting protocol, neither signal was privileged; tuning target-specific coefficients against retrospective recovery would have introduced outcome-dependent selection. The equal weighting was fixed across targets, methods, and seeds and did not imply biological equivalence, statistical independence, equal predictive value, or optimal weighting.

The resulting *O_i_* was a count-aware, multi-round NGS evidence target rather than an apparent-affinity estimate. It served as the common target for global pairwise fitting and, within AbPACER, as a source of neighbor evidence. Final prioritization used each method’s learned output score; AbPACER candidates were ranked by *s_i_*, not directly by *O_i_*.

Deterministic controls included total and round-specific read counts; *Z*_2_*_/_*_1_, *Z*_3_*_/_*_1_, *Z*_3_*_/_*_2_, and *T*_peak_; direct *O_i_* ranking; count–*O_i_* rank fusion; and *O_i_*ranking within exact total-count ties. No control was selected using apparent-affinity values. Complete definitions, tie handling, and recovery profiles are provided in Additional file 1.

### 2.4 Parent-aware representation and two-stage neural ranking

Affinity-maturation candidates were represented as changes from their target-specific parental clone rather than as absolute sequences. This representation matched the experimental setting because each candidate library was generated by mutating a known parental sequence, and the interpretation of a substitution depends on its chain, position, and sequence background [15, 16]. Differences in VH and VL were identified separately while preserving paired-chain identity. Each mutation token encoded chain identity, an approximate parent-referenced framework/CDR region, one-based model position, parental and candidate residues, and standardized physicochemical changes. Region labels were modeling features derived from parent-referenced raw positions rather than curated structural or IMGT annotations.

Explicit mutation descriptors retained the exact and auditable identity of each parent-to-candidate change, whereas frozen IgBERT provided sequence context not represented by the hand-defined fields. Parental and candidate sequences were encoded separately in VH [SEP] VL format. At each differing position, the candidate and parental residue embeddings, their signed difference, and their absolute difference were merged with the explicit mutation descriptor. The signed difference retained the direction of the parent-to-candidate contextual shift, whereas the absolute difference retained its magnitude independently of direction. A separate global token summarized the full paired-sequence shift and complemented the residue-level mutation tokens.

A two-layer, four-head transformer jointly processed the global token and the variable-length set of context-aware mutation tokens, allowing information from mutations distributed across VH and VL to interact instead of treating each mutation as an independent contribution. The global-token output provided a fixed-length query representation *q_i_*. IgBERT remained frozen so that pretrained antibody-sequence context could be used without adapting the full language model to the noisy affinity-label-blind NGS target. Task-specific learning was therefore restricted to the mutation encoder, context projections, merge network, transformer, and scoring modules. IgBERT functioned as a contextual feature extractor rather than a pretrained affinity predictor.

### Neighbor-aware base scoring

The NGS evidence associated with an individual clone can be sparse or noisy, whereas unmeasured clones undergoing related parent-relative sequence changes may provide additional within-campaign evidence. Base neighbors were therefore retrieved from an unmeasured reference pool by cosine similarity in normalized signed candidate-minus-parent IgBERT mean-difference space. Using a parent-relative difference space aligned retrieval with the affinity-maturation process, and cosine similarity emphasized the direction of the contextual shift while reducing dependence on embedding magnitude. Retrieval similarity was used only to identify potentially informative reference clones and was not treated as evidence that nearby sequences had the same molecular affinity.

Quantitatively measured clone keys were excluded from the reference pool, so neighbor aggregation used only affinity-label-blind *O_j_*values from unmeasured clones. Up to 32 neighbors were fixed for each query as a computational cap rather than a biological neighborhood definition. Because not every retrieved clone was expected to be equally informative, query-conditioned attention combined relation features with neighbor *O_j_* evidence instead of applying uniform averaging. A learned null-neighbor option allowed the model to reject an uninformative retrieval set rather than being forced to borrow evidence from weakly related clones. Auxiliary neighborhood summaries *a_i_* exposed the strength and reliability of the retrieved support alongside the attention-weighted neighbor representation *n_i_*. The combined feature was

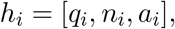

which a multilayer perceptron mapped to the neighbor-aware base score *b_i_*.

### Global pairwise training

The base stage learned the relative ordering of *O_i_* through a weighted pairwise softplus objective. Pairs with larger *O_i_* separation and stronger NGS support received greater emphasis because their relative ordering was better resolved by the affinity-label-blind evidence. No pointwise regression term was used. Pairwise learning was aligned with the downstream prioritization task and avoided treating absolute differences in *O_i_* as calibrated molecular or biological distances.

Checkpoints were selected by Spearman correlation with *O_i_* on unmeasured validation rows. After the training duration had been selected, a newly initialized final base model was fitted for that number of epochs using all unmeasured rows. This final refit used the available affinity-label-blind training evidence without using the retrospective measured panel, and the resulting base model was frozen before local refinement.

### Bounded local-family refinement

The base stage learned a broad ordering across the candidate universe, whereas the practical assay decision required finer discrimination among already count-supported candidates near the primary screening region. The residual stage was therefore restricted to the pre-exclusion top-5% count-supported region. Its membership was fixed from total count before measured-clone exclusion and recovery evaluation, so the local stage did not define the candidate universe from retrospective outcomes.

The local-family graph was separate from the base-attention retrieval graph and combined three complementary relations: frozen-IgBERT parent-delta similarity, total-count similarity, and exact mutation overlap. Combining parent-delta context and count similarity with an upweighting for exact mutation overlap made the local relation more conservative than relying on sequence similarity alone. Relation-weighted neighbor *O_j_* values supplied the local ordering signal, and winsorization limited the influence of extreme neighbor targets.

A zero-initialized residual head received *h_i_* and produced

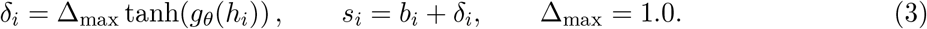

The zero initialization made the initial refined score equal to the base score, *s_i_* = *b_i_*. Base parameters remained frozen during residual fitting, and the hyperbolic-tangent bound limited any local correction to one score unit. Together with the separate local graph, these constraints allowed the second stage to refine local ordering without relearning or replacing the global ranker.

Residual checkpoints were selected by held-out local anchor-pair loss. A newly initialized residual head was then fitted for the selected number of epochs using all eligible unmeasured local-family rows while the base model remained frozen. The final AbPACER score was *s_i_*; candidates were ordered by decreasing *s_i_*, with exact ties resolved by the frozen paired VH|VL key. Exact feature dimensions, attention inputs, pairwise losses, local-relation formulas, retrieval caps, winsorization, and edge-case handling are provided in Additional file 1.

### 2.5 Common-protocol comparator adaptations and controlled ablations

Learned comparators were evaluated as common-protocol adaptations or reimplementations rather than exact reproductions of their original tasks. They comprised an Ens-Grad-inspired equal-weight ensemble of six one-hot one-dimensional convolutional neural network (CNN) models [9], end-to-end ESM-2-based AbAffinity [21, 13], and PALM-based A2Binder [22]. For the phage-display analysis, A2Binder used only the released heavy-and light-chain branches because its 300-token antigen pathway could not accommodate the complete antigen sequences without substantial truncation; this adaptation is denoted A2Binder-HL.

Within each target and seed, all learned pipelines used identical training and validation clone keys, the same affinity-label-blind target *O_i_*, presampled global-pair schedules, the same weighted global pairwise objective, frozen candidate manifests, seeds 42–44, no calibration, and a common deterministic score-tie rule. Comparator models and the AbPACER base stage used validation-Spearman checkpoint selection, whereas AbPACER additionally applied bounded local residual refinement. Architecture-specific parameterization and optimization remained part of the complete-pipeline comparison.

Controlled ablations removed query-level IgBERT context, local residual refinement, or neighbor-derived forward features while retaining the common protocol. The no-query-context condition retained explicit mutation descriptors, the IgBERT-derived neighbor graph, and local-family supervision; the no-residual condition ranked candidates by *b_i_*; and the no-neighbor-forward condition removed *n_i_* and *a_i_* while retaining the local-family graph and residual supervision. Because these interventions changed trainable parameter counts, they were interpreted as pathway-level probes rather than parameter-matched component tests. Exact adaptations, ablation definitions, parameter accounting, and training audits are provided in Additional file 1.

### 2.6 AlphaSeq common-split benchmark

The AlphaSeq arm evaluated supervised adaptation of the parent-aware representation separately from the phage-display workflow. The AbAffinity-processed cohort comprised 71,834 variants across the 14H, 14L, 91H, and 95L libraries, with fixed fit, validation, and test assignments of 51,139, 9,025, and 11,670 rows [12, 13, 23]. All methods used these assignments and seeds 42–44, optimized mean-squared error or an equivalent pointwise squared-error objective, selected checkpoints by validation Pearson correlation, and produced uncalibrated test predictions. Test labels were accessed only after predictions had been frozen for final evaluation. This benchmark used no phage-display R1–R3 counts or trajectories, phage target *O_i_*, or phage NGS-derived neighbor evidence.

One shared AbPACER-MSE model was trained per seed across all four libraries to test whether parent-relative adaptation could be learned through a single shared set of task-specific parameters. Each variant retained its library-specific parental and candidate sequences, parent-relative features, frozen IgBERT context, and within-library feature and neighbor construction; no library-identity embedding was used. Neighbor references were restricted to fit-set rows from the corresponding library. Within each library, bandwidth was selected by validation Spearman correlation using support calculated only from fit-set neighbors; validation and test rows were not used as neighbor references.

With *K_D_*denoting the equilibrium dissociation constant and *y_i_* = log_10_(*K_D,i_*), targets were sign-reversed so that larger normalized values represented stronger binding and were standardized using pooled fit-cohort statistics. The fit-derived transformation was applied unchanged to validation and test rows. The base and bounded residual heads were trained sequentially by mean-squared error, with the base frozen during residual fitting, and predictions were inverse-transformed before evaluation. AbAffinity and A2Binder each used one pooled model per seed, whereas Ens-Grad CNN averaged six pooled one-hot Conv1D regressors. AbAffinity fine-tuned ESM-2 t33 650M end to end, and AlphaSeq A2Binder retained its heavy-, light-, and antigen-sequence branches. Parameter counts included all unique task-specific parameters updated per seed across the deployed model or ensemble constituents and excluded frozen or non-trainable components. Exact target transformations, comparator adaptations, and parameter accounting are provided in Additional file 1.

The primary fixed-budget endpoint ranked all 11,670 frozen test predictions and selected 384 variants, matching the 384-candidate selection depth used in the phage-display arm. The measured comparison set comprised the 384 test variants with the lowest *K_D_* values, and fixed-budget recovery was the overlap between the predicted and measured top-384 sets. Label or prediction ties at the selection boundary were resolved using the deterministic fixed-test row key. Global Pearson correlation, Spearman correlation, root-mean-square error (RMSE), mean absolute error (MAE), and within-library pairwise metrics provided complementary assessments of prediction and ranking behavior.

To reduce the influence of library size, between-library difficulty, and cross-library prediction-scale differences, a complementary analysis reused the frozen pooled predictions and calculated metrics separately within each library before macro-averaging. These metrics were 10×-confident area under the receiver operating characteristic curve (ROC–AUC) for pairs separated by at least one log_10_(*K_D_*) unit, all-pair concordance with half credit for predicted ties, and Kendall’s *τ_b_*. These within-library ranking metrics were based on an AbRank-inspired evaluation principle but used neither the AbRank dataset nor its splits [24]. Four separately trained library-specific AbPACER-MSE models were retained as a training-granularity sensitivity analysis in Additional file 1.

### Use of generative AI-assisted tools

The authors were responsible for the scientific conception, study design, experimental work, analyses, interpretation, and original manuscript content. OpenAI ChatGPT and Prism assisted with English-language editing, readability, and organization of author-written text, and OpenAI Codex assisted with code implementation, debugging, refactoring, and testing under author-defined specifications. All AI-assisted outputs were reviewed and, where applicable, tested and verified by the authors, who take responsibility for the final manuscript, software, analyses, and conclusions.

## 3 Results

### 3.1 Primary fixed-budget phage-display prioritization

Both full pools contained extensive low-count tails, and the frozen top-5% candidate sets still contained 16,323 FAF1 and 7,487 VEGFR candidates, far exceeding the 384-candidate selection depth. Count resolution was particularly coarse in FAF1, where 4,626 candidates shared total count 6; the largest VEGFR exact-count tie comprised 359 candidates at count 19.

All clones in the fixed retrospective evaluation panels had lower target-specific ELISA-derived apparent-affinity values than their respective parental clones.

The uniform-random expectation was 0.165 recovered clones in FAF1 and 0.154 in VEGFR. Total-count ranking substantially exceeded this reference in both campaigns, recovering one of seven FAF1 panel clones and all three VEGFR panel clones. Total count therefore provided a strong operational baseline rather than a trivial comparator.

At the primary 384-candidate endpoint, AbPACER achieved the highest observed mean recovery among the learned methods in both campaigns. In FAF1, AbPACER recovered two panel clones in each of the three seeds, compared with one by total count. It was also the only learned method to recover two FAF1 panel clones consistently across all seeds.

VEGFR showed a different ranking regime. AbPACER recovered 2.33 ± 0.58 of three panel clones and achieved the highest mean among the learned methods, whereas total count already placed all three panel clones within the top 384. Thus, sequence-conditioned reranking provided its clearest improvement over count in FAF1, while the deterministic count baseline remained strongest in VEGFR.

The recovered panel members therefore provide retrospective evidence that campaign-specific 384-candidate lists can contain previously measured clones with favorable apparent-affinity changes relative to their parental clones. Because apparent affinity was unavailable for the remaining selected candidates, these recoveries do not estimate the fraction of favorable clones or the precision of the complete 384-candidate lists.

Broader hypothetical budgets are reported in Additional file 1. No learned method was uniformly best across those depths, and AbPACER’s clearest separation from the learned comparators occurred at the primary 384-candidate endpoint. Direct *O_i_* ranking and the deterministic count and trajectory controls did not reproduce the FAF1 top-384 recovery, indicating that the result was not obtained by directly ordering the common scalar supervision target.

Model size alone also did not explain the primary result. AbPACER used 1.38 million task-specific trainable parameters, compared with 114.63 million for A2Binder-HL and 651.04 million for AbAffinity, whereas the smaller Ens-Grad CNN showed lower top-384 recovery.

### Candidate-level interpretation of the FAF1 gain

The additional FAF1 panel clone recovered by AbPACER at the 384-candidate endpoint was FAF1_#138, which had an ELISA-derived apparent affinity value of 27.8 pM compared with 43.1 pM for parental FAF1_#1. Its total count of 6 placed it in a 4,626-candidate exact-count tie spanning ranks 11,698–16,323. Thus, even the most favorable ordering within that tie could place the clone no higher than rank 11,698, far outside the assay budget. AbPACER instead ranked FAF1_#138 at 361, 317, and 359 across seeds 42–44, respectively (Table 3). The one-clone recovery difference therefore corresponded to a concrete assay-list decision: a measured low-count clone that no total-count-preserving tie-break could reach was included in every frozen AbPACER top-384 list.

**Table 3:**
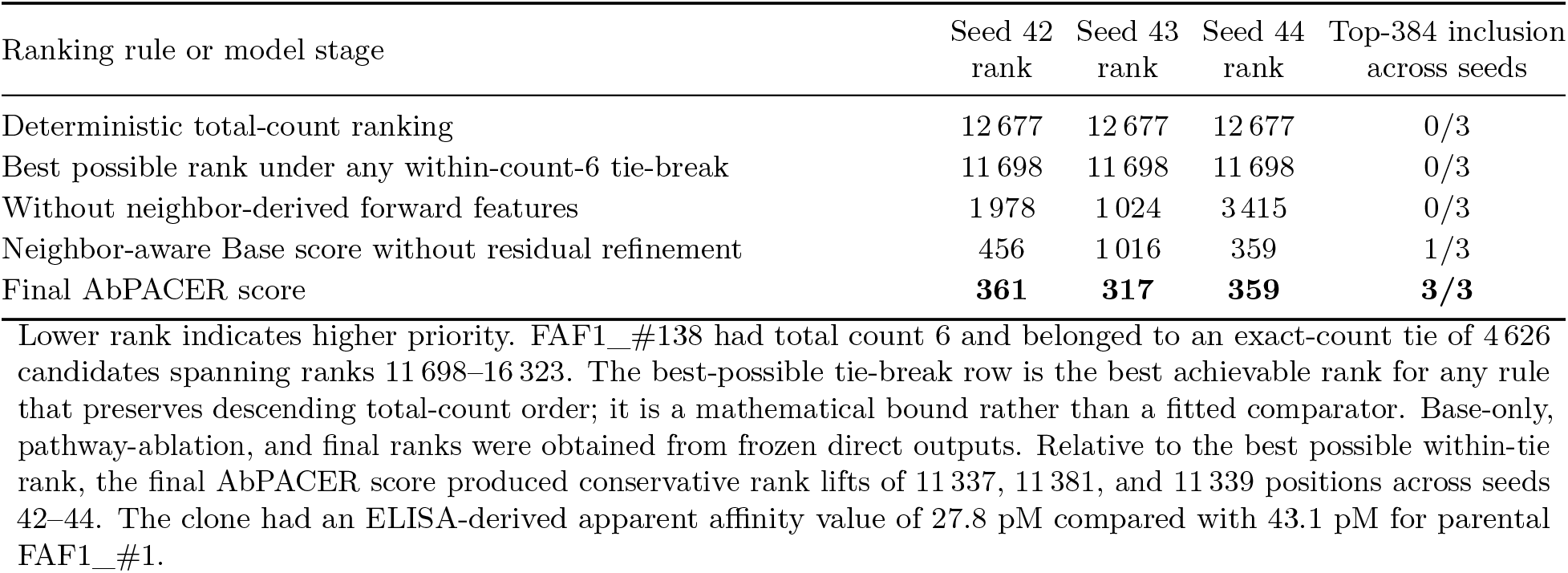
Tie-aware and stage-specific rank audit of FAF1_#138 at the primary 384-candidate assay boundary.

| Ranking rule or model stage | Seed 42<br>rank | Seed 43<br>rank | Seed 44<br>rank | Top-384 inclusion<br>across seeds |
| --- | --- | --- | --- | --- |
| Deterministic total-count ranking | 12 677 | 12 677 | 12 677 | 0/3 |
| Best possible rank under any within-count-6 tie-break | 11 698 | 11 698 | 11 698 | 0/3 |
| Without neighbor-derived forward features | 1 978 | 1 024 | 3 415 | 0/3 |
| Neighbor-aware Base score without residual refinement | 456 | 1 016 | 359 | 1/3 |
| Final AbPACER score | **361** | **317** | **359** | **3/3** |

Lower rank indicates higher priority. FAF1\_#138 had total count 6 and belonged to an exact-count tie of 4626 candidates spanning ranks 11 698–16 323. The best-possible tie-break row is the best achievable rank for any rule that preserves descending total-count order; it is a mathematical bound rather than a fitted comparator. Base-only, pathway-ablation, and final ranks were obtained from frozen direct outputs. Relative to the best possible within-tie rank, the final AbPACER score produced conservative rank lifts of 11 337, 11 381, and 11 339 positions across seeds 42–44. The clone had an ELISA-derived apparent affinity value of 27.8 pM compared with 43.1 pM for parental FAF1\_#1.

**Table 4:** Common-split AlphaSeq benchmark at the shared 384-selection depth.

**A. Global fixed-budget top-384 overlap**
| Model | Task-specific trainable parameters | True top-384 variants recovered among 384 selected |
| --- | --- | --- |
| Uniform random selection (expected) | – | 12.64 |
| AbPACER-MSE | 1.378 M | $187.0 \pm 2.6$ |
| AbAffinity | 651.04 M | **$188.0 \pm 2.6$** |
| A2Binder | 263.85 M | $175.7 \pm 6.4$ |
| Ens-Grad CNN | 0.058 M | $154.0 \pm 6.1$ |

**B. Global regression performance**
| Model | Task-specific trainable parameters | Pearson<br>↑ | Spearman<br>↑ | RMSE<br>↓ | MAE<br>↓ |
| --- | --- | --- | --- | --- | --- |
| AbPACER-MSE | 1.378 M | **$0.687 \pm 0.003$** | **$0.652 \pm 0.002$** | **$0.973 \pm 0.004$** | **$0.741 \pm 0.003$** |
| AbAffinity | 651.04 M | $0.686 \pm 0.002$ | **$0.652 \pm 0.001$** | $0.975 \pm 0.003$ | $0.743 \pm 0.002$ |
| A2Binder | 263.85 M | $0.676 \pm 0.003$ | $0.645 \pm 0.003$ | $1.006 \pm 0.019$ | $0.759 \pm 0.010$ |
| Ens-Grad CNN | 0.058 M | $0.634 \pm 0.003$ | $0.604 \pm 0.003$ | $1.088 \pm 0.002$ | $0.879 \pm 0.002$ |

**C. Within-library pairwise ranking**
| Model | Task-specific trainable parameters | 10×-confident AUC ↑ | All-pair concordance ↑ | Kendall $\tau_b$ ↑ |
| --- | --- | --- | --- | --- |
| AbPACER-MSE | 1.378 M | **$0.877 \pm 0.002$** | $0.694 \pm 0.001$ | $0.388 \pm 0.002$ |
| AbAffinity | 651.04 M | **$0.877 \pm 0.001$** | **$0.695 \pm 0.000$** | **$0.394 \pm 0.001$** |
| A2Binder | 263.85 M | $0.874 \pm 0.001$ | $0.692 \pm 0.000$ | $0.385 \pm 0.001$ |
| Ens-Grad CNN | 0.058 M | $0.847 \pm 0.001$ | $0.677 \pm 0.001$ | $0.355 \pm 0.001$ |

Values for learned methods are mean ± sample standard deviation over seeds 42–44. Every method ranked the same fixed test set of 11 670 variants and selected its predicted top 384. The oracle set comprised the 384 variants with the lowest measured $K_D$ values. Because both predicted and oracle sets contained 384 variants, precision@384 and recall@384 were numerically identical. The expected uniform-random overlap was $384 \times 384 / 11\,670 = 12.64$ . Parameter counts are per-seed totals of unique task-specific parameters updated across all deployed models or ensemble constituents and exclude frozen or non-trainable components. Panel C metrics were calculated separately within each library and macro-averaged across the four libraries. The 10×-confident AUC used only pairs separated by at least one $\log_{10}(K_D)$ unit. All-pair concordance included all within-library pairs and assigned half credit to predicted ties. Kendall’s $\tau_b$ measured complete rank agreement while accounting for ties. These metrics followed an AbRank-inspired evaluation principle but used neither the AbRank dataset nor its splits.
Boldface denotes the numerically highest observed mean within each metric; rounded ties are jointly bolded. Formatting does not imply statistical significance.

The stage-specific ranks separated global promotion from boundary refinement. The neighbor-aware Base score had already moved FAF1_#138 from deterministic count rank 12,677 to ranks 456, 1,016, and 359 across seeds 42–44. The large global promotion therefore occurred before local residual refinement. The bounded residual subsequently changed these ranks to 361, 317, and 359, converting Base-stage top-384 inclusion in one of three seeds into final inclusion in all three seeds. For this clone, the Base stage supplied the global sequence-conditioned promotion, whereas bounded residual refinement stabilized placement at the 384-candidate assay boundary.

At the aggregate recovery level, removing neighbor-derived forward features reduced FAF1 top-384 recovery from 2.00 ± 0.00*/*7 to 1.00 ± 0.00*/*7 and VEGFR recovery from 2.33 ± 0.58*/*3 to 0.67 ± 0.58*/*3. Removing local residual refinement reduced FAF1 recovery to 1.33 ± 0.58*/*7 while leaving VEGFR recovery unchanged. The complete verified representation and scoring/refinement ablations, including the target-dependent query-level IgBERT result, are reported in Additional file 1. Because the ablated models were not parameter matched, these comparisons support pathway-level interpretation rather than component-wise causal attribution.

### Secondary selection-pattern analysis

A parent-referenced VH-position-57 F→L/I motif defined before panel annotation showed the same direction of enrichment in all three seeds when FAF1 candidates selected only by AbPACER were compared with total-count top-384 candidates displaced from the corresponding AbPACER list. The motif occurred in 59.0%, 32.4%, and 52.1% of the measured-panel-excluded AbPACER-only selections across seeds 42–44, respectively, compared with 1.3%, 1.9%, and 1.4% of the corresponding count-only displaced candidates. Evaluation-panel clones were excluded from feature discovery, statistical testing, support filtering, and stability assessment. After the mutation-analysis feature set had been frozen, panel annotation showed that the same motif was present in five of the seven FAF1 evaluation-panel clones, including FAF1_#138. This concordance identifies a recurrent target-specific signature of the resulting AbPACER allocation. Because the compared groups differed substantially in their count-rank distributions and apparent affinity was unavailable for the unmeasured candidates, the analysis was not interpreted as a count-independent mutation-effect test, evidence of a causal affinity effect, or proof that all AbPACER-only candidates were superior.

### 3.2 Supervised AlphaSeq adaptation at the shared 384-selection depth

The AlphaSeq arm tested whether the parent-aware representation could support affinity ranking in a separate densely labeled setting without phage-display R1–R3 counts or trajectories, the phage-specific target *O_i_*, or phage NGS-derived neighbor evidence. At the primary endpoint, which used the same 384-candidate budget as the phage-display arm, AbPACER-MSE recovered 187.0 ± 2.6 of the true top-384 variants among 384 selected predictions, compared with 188.0 ± 2.6 for AbAffinity, 175.7 ± 6.4 for A2Binder, and 154.0 ± 6.1 for Ens-Grad CNN. Seed-specific overlaps were 188, 189, and 184 for AbPACER-MSE and 190, 189, and 185 for AbAffinity, corresponding to paired differences of −2, 0, and −1 variants. Thus, the one-variant difference between their aggregate means reflected consistent near-parity across seeds rather than compensation between highly divergent runs.

Global regression performance showed the same pattern. AbPACER-MSE achieved Pearson 0.687 ± 0.003, Spearman 0.652 ± 0.002, RMSE 0.973 ± 0.004, and MAE 0.741 ± 0.003, compared with 0.686 ± 0.002, 0.652 ± 0.001, 0.975 ± 0.003, and 0.743 ± 0.002, respectively, for AbAffinity. Because pooled regression metrics can be influenced by library size, between-library difficulty, and cross-library prediction-scale differences, we additionally evaluated mutation-variant ordering separately within each AlphaSeq library and macro-averaged the results across libraries.

Following an AbRank-inspired confident-pair evaluation principle [24], the 10×-confident AUC retained only within-library variant pairs separated by at least one log_10_(*K_D_*) unit, corresponding to a minimum tenfold difference in measured affinity. It therefore assessed whether a model correctly distinguished the stronger-binding member of clearly separated variant pairs, while reducing the influence of near-ties. All-pair concordance evaluated ordering across every within-library pair and assigned half credit to predicted ties, providing a broad measure of pairwise ranking consistency. Kendall’s *τ_b_* quantified agreement between the complete predicted and measured within-library rankings while accounting for ties. These three metrics assessed confident affinity separation, general pairwise ordering, and overall ranked-list agreement. The analysis followed an AbRank-inspired evaluation principle but used neither the AbRank dataset nor its splits.

AbPACER-MSE and AbAffinity both achieved a 10×-confident AUC of 0.877. Their all-pair concordances were 0.694±0.001 and 0.695±0.000, and their Kendall’s *τ_b_* values were 0.388±0.002 and 0.394 ± 0.001, respectively. AbPACER-MSE therefore closely matched AbAffinity not only in global affinity regression and fixed-budget top-384 recovery, but also in ordering parent-related mutation variants within individual libraries. A2Binder was numerically close on the three within-library metrics, whereas Ens-Grad CNN was consistently lower.

Pooled AbPACER-MSE updated 1.378 million task-specific parameters, approximately 472-fold fewer than AbAffinity and 191-fold fewer than A2Binder. The lightweight Ens-Grad CNN updated fewer parameters than AbPACER-MSE but showed lower fixed-budget recovery, global regression, and within-library ranking performance, indicating that parameter count alone did not explain the observed result.

Training-granularity sensitivity showed essentially identical global regression and within-library pairwise-ranking performance for pooled and four-library-specific AbPACER-MSE. The pooled model used one quarter of the task-specific trainable parameters while retaining the correct library-specific parental sequence for each variant and using no explicit library-identity embedding. Secondary threshold-based recovery values are reported in Additional file 1. This result indicates that separate library-specific models were not required to retain the observed regression and ranking performance.

Across these endpoints, AbPACER-MSE retained near-AbAffinity performance while updating approximately 472-fold fewer task-specific parameters, supporting efficient supervised adaptation of the parent-aware representation on a separate densely labelled affinity landscape.

## 4 Discussion

This study evaluated AbPACER as a campaign-adaptive fixed-budget prioritizer rather than as a calibrated affinity predictor or a model transferred unchanged across campaigns. The two retrospective phage-display campaigns represented distinct operational regimes. In FAF1, coarse count resolution and a large exact-count tie left room for sequence-conditioned reranking, and AbPACER recovered two of seven panel clones in every seed compared with one by total count. In VEGFR, total count already recovered all three panel clones, whereas AbPACER achieved the highest learned-method mean without exceeding that deterministic ceiling. The phage results therefore support a targeted benefit at the 384-candidate assay boundary under count ambiguity, rather than uniform superiority across targets or broader ranking depths.

The FAF1 improvement was numerically narrow but operationally concrete. FAF1_#138 belonged to a count-6 tie whose best possible count-preserving rank was 11,698, so its inclusion in all three AbPACER top-384 lists could not be explained by arbitrary ordering within an exact-count tie. Direct ranking by the common affinity-label-blind target *O_i_*also left the clone outside the assay boundary. The neighbor-aware Base score supplied the large global promotion, whereas bounded residual refinement converted Base-stage inclusion in one seed into final inclusion in all three. Consistent with this stage audit, removing neighbor-derived forward features reduced top-384 recovery in both targets, whereas removal of residual refinement mainly affected FAF1. Because the ablations were not parameter matched and the evaluation panels were small, these comparisons support pathway-level interpretation rather than component-wise causal attribution.

An analysis of frozen outputs that excluded measured panel clones identified a recurrent FAF1 selection pattern. A parent-referenced VH-position-57 F→L/I motif was repeatedly enriched among candidates selected only by AbPACER and, after the mutation-analysis feature set had been frozen, was observed in five of the seven retrospective FAF1 panel clones. This agreement indicates that the resulting allocation was associated with a recurrent parent-relative mutation pattern rather than an arbitrary rearrangement of candidate identities. However, the AbPACER-only and count-only displaced groups differed substantially in count-rank distribution, and apparent-affinity measurements were unavailable for the unmeasured candidates. The analysis therefore does not establish a count-independent mutation effect, a causal role for VH position 57, or superior affinity among the remaining AbPACER-only selections.

AlphaSeq provided complementary evidence under a distinct supervision regime without phage-display R1–R3 counts or trajectories, the phage-specific target *O_i_*, or phage NGS-derived neighbor evidence. AbPACER-MSE recovered 187.0±2.6 of the true top-384 variants compared with 188.0±2.6 for AbAffinity and remained within 0–2 recovered variants of AbAffinity in every seed. Their global regression metrics were nearly identical, and the agreement extended to three complementary within-library ranking assessments: discrimination of variant pairs with at least tenfold affinity separation, ordering across all variant pairs, and agreement between the complete predicted and measured rankings. AbPACER-MSE closely matched AbAffinity across these measures, although A2Binder was also numerically close in the within-library analysis. AbPACER-MSE achieved this performance while updating approximately 472-fold fewer task-specific parameters than AbAffinity and 191-fold fewer than A2Binder. Moreover, one pooled model retained essentially the same regression and within-library ranking performance as four library-specific models while using one quarter of the task-specific parameters and no explicit library-identity embedding. These comparisons do not establish computational efficiency or cross-antigen validation of the complete phage-display workflow.

AbPACER is intended for campaign-adaptive use: it uses campaign-specific sequence and R1–R3 evidence to order the frozen count-supported candidate universe, while total count remains an empirical-support filter and deterministic baseline. The output is a prioritization ranking, not a calibrated affinity estimate. The 384-candidate endpoint is neither a biologically optimal nor universal assay size; laboratories may truncate the complete ranking to match their downstream screening capacity. The primary claims here are specific to the reported endpoint.

The principal limitation of the phage-display evaluation is its retrospective design and the small quantitative panels of seven FAF1 and three VEGFR clones. Because the remaining selected candidates were unmeasured, the study cannot estimate list-level precision, affinity distributions, or generality across campaigns. Prospective validation was not feasible within the present study because additional affinity-maturation campaigns and quantitative characterization of a sufficiently large candidate panel would require substantial experimental time and resources. Future studies should freeze the candidate manifest, model configuration, seeds, and assay list before measurement and evaluate larger panels across additional targets. The planned release of paired-chain data, predictions, complete rankings, and code will support independent reanalysis and such follow-up studies.

## 5 Conclusions

AbPACER ranks observed paired scFv candidates using parent-relative sequence features and affinity-label-blind phage-display NGS evidence. At the 384-candidate endpoint, it showed the highest mean recovery among learned methods in both retrospective campaigns. In FAF1, it consistently recovered one additional measured clone that no count-preserving ranking could place within the assay budget; in VEGFR, total count remained strongest. On AlphaSeq, pooled AbPACER-MSE closely matched AbAffinity across top-384 recovery, global regression, and within-library ranking while updating approximately 472-fold fewer task-specific parameters. Prospective frozen-list evaluation across additional targets and larger quantitative panels is required to determine list-level performance and generality.

## Additional files

**Additional file 1 (.pdf).** Supplementary methods and implementation details, frozen candidate manifests, retrospective measurement and rank audits, deterministic controls, completed representation and scoring/refinement ablations, tie-aware clone-level audits, measured-panel-excluded mutation-pattern analyses, comparator adaptations and task-specific parameter accounting, AlphaSeq seed-, library-, and training-granularity analyses, provenance, and validation-based training audits.

## Declarations

### Ethics approval and consent to participate

Not applicable.

### Consent for publication

Not applicable.

### Availability of data and materials

Public AlphaSeq data are available from the original Zenodo record [23]. The original AlphaSeq sequence and measured-label cohort will not be redistributed. The AbPACER release will provide label-stripped predictions and derived benchmark outputs, together with provenance identifiers or hashes and a citation and link to the upstream record.

Raw PacBio reads from the FAF1 and VEGFR affinity-maturation campaigns are associated with NCBI BioProject PRJNA1506661 and will be released publicly through the NCBI Sequence Read Archive no later than publication. Processed phage-display datasets are being archived in Zenodo and will include paired VH/VL sequences, R1–R3 and total read counts, parent-relative annotations, frozen candidate-manifest memberships, and retrospective ELISA-derived apparent affinity metadata.

Versioned code and reproducibility artifacts are hosted in the AbPACER GitHub repository [25] and will be made public and archived in Zenodo no later than publication. These artifacts will include resolved configurations, train/validation split memberships, frozen candidate-manifest memberships, candidate-level predictions, complete direct rankings, frozen top-384 assay lists, evaluation outputs, software-environment records, and checksums or hashes. The software will be released under the BSD-3-Clause license, and the phage-display data and article-associated artifacts under CC BY 4.0. AngioLab, Inc., which generated the experimental datasets, authorized their public release.

## Funding

AngioLab, Inc. provided financial and in-kind support for the experimental component of this study, including the phage-display affinity-maturation experiments and binding ELISA measurements, and supported the PacBio Revio sequencing. AngioLab, Inc. contributed to experimental data generation and curation through BYP and EBP and authorized the public release of the experimental datasets. AngioLab, Inc. had no role in the design or execution of the computational study, development or evaluation of AbPACER, computational analyses, interpretation of the computational results, or preparation of the original manuscript.

JHH was supported in part by the MSIT (Ministry of Science and ICT), Korea, under the ITRC (Information Technology Research Center) Support Program (IITP-2026-RS-2022-00156225) supervised by the IITP (Institute for Information and Communications Technology Planning and Evaluation), and in part by the National Research Foundation of Korea (NRF) grant funded by the Korean government (No. RS-2024-00415812).

### Authors’ contributions

AJC conceived the computational study; designed, developed, and evaluated AbPACER; performed the computational analyses; interpreted the results; and wrote the original draft of the manuscript. BYP and EBP performed the phage-display affinity-maturation experiments and generated and curated the PacBio-derived and binding ELISA datasets. JHH supervised the doctoral research, reviewed the study design and interpretation, critically revised the manuscript, and approved the final version. All authors read and approved the final manuscript.

## Acknowledgements

The authors thank AngioLab, Inc. for authorizing the public release of the experimental datasets and associated metadata.

## List of abbreviations

AbPACER: Antibody Parent-Aware Contextual Evidence Ranker
AUC: area under the receiver operating characteristic curve
CDR: complementarity-determining region
CNN: convolutional neural network
ELISA: enzyme-linked immunosorbent assay
FAF1: Fas-associated factor 1
*K_D_*: equilibrium dissociation constant
MAE: mean absolute error
MSE: mean-squared error
NGS: next-generation sequencing
RMSE: root-mean-square error
scFv: single-chain variable fragment
scFv–Fc: single-chain variable fragment–Fc fusion
VEGFR: vascular endothelial growth factor receptor
VH: variable heavy chain
VL: variable light chain.

