## Additional file 1: Supplementary methods, figures, and tables for "AbPACER: parent-aware, affinity-label-blind prioritization of affinity-matured scFv clones from phage-display NGS"

#### Supplementary information overview

AbPACER denotes the Antibody Parent-Aware Contextual Evidence Ranker, and AbPACER-MSE denotes its supervised AlphaSeq adaptation. The frozen top-5% candidate manifests define the primary phage-display analyses. The top-1% and top-10% manifests are retained only as narrower and broader reference definitions for documenting the scale and panel coverage surrounding the primary operational choice. All learned primary comparisons use direct frozen model outputs, fixed seeds 42, 43, and 44, fixed train/validation memberships, and deterministic score-tie handling. Across the affinity-label-blind phage and supervised AlphaSeq arms, the primary fixed-budget endpoint uses a selection depth of 384 as an operational high-throughput downstream-screening benchmark motivated by 384-well antibody-screening and immunoassay formats. Comparator labels refer to common-protocol adaptations or reimplementations rather than exact reproductions of the original tasks. AbPACER is intended to be fitted separately for each affinity-maturation campaign using the available paired sequences and round-resolved R1–R3 counts before producing a direct 384-candidate assay list.

This document provides the implementation details and aggregate audits required to interpret the study, including preprocessing, fitting, direct ranking, ablations, comparator adaptations, AlphaSeq analyses, provenance, and a validation-based training-sufficiency and checkpoint audit. The complete machine-readable reproducibility record will be provided in the versioned archival release rather than duplicated in this PDF. Section 1 follows the methodological sequence of the main manuscript. Section 5 contains completed component ablations, tie-aware clone-level rank audits, and measured-panel-excluded mutation-pattern analyses; the remaining sections provide cohort audits, comparator adaptations and parameter accounting, benchmark results, provenance records, and the validation-based training audit.

**2 Full-pool statistics, frozen candidate manifests, and retrospective measurement audit** **10**

**3 Parent-to-candidate mutation maps** **13**

**4 Deterministic controls, direct learned rankings, and clone-level audit** **14**

**5 Representation, scoring/refinement, and frozen-output candidate-level analyses** **16**

**6 Comparator adaptations and task-specific parameter accounting** **18**

**7 Detailed AlphaSeq results and sensitivity analyses** **19**

**8 Manifest provenance, cohort definition, and release scope** **21**

**9 Validation-based training-sufficiency and checkpoint audit** **22**

### 1 Supplementary methods and implementation details

#### 1.1 Experimental generation and sequence preprocessing

Parental scFv genes in pDR-D1 were diversified with the GeneMorph II Random Mutagenesis Kit under low-template conditions. The VEGFR clone #163 library was generated using four successive mutagenic PCR amplifications. For FAF1_#1, pilot libraries were evaluated for deletion and stop-codon frequency, and the final production library was generated from the third-round PCR product selected during library optimization. The final inserts were SfiI-digested, ligated into pDR-D1, electroporated into electrocompetent *E. coli* XL1-Blue cells, recovered in SOC medium, and stored as target-specific affinity-maturation library stocks. Library phage was rescued with VCSM13 helper phage in *E. coli* ER2738 and precipitated with PEG8000/NaCl.

The target-specific affinity-maturation libraries underwent three rounds of solid-phase panning in 96-well microplates with 4% skim-milk blocking, progressively lower antigen concentrations, and target-specific wash schedules. Antigens were coated overnight at 10, 1, and 0.1 nM for FAF1 and at 9, 0.9, and 0.09 nM for VEGFR-2 across R1–R3. For FAF1, 20, 30, and 40 washes were performed in R1, R2, and R3, with wash/contact times of 1, 2, and 3 min, respectively. For VEGFR-2, R1 comprised 30 washes without a dwell interval followed by 10 washes with a 1-min interval; R2 and R3 each comprised 40 washes with 1- and 3-min intervals, respectively. Washing used buffer containing 0.05% Tween-20. Bound phage was sequentially eluted with glycine–HCl and triethylamine, neutralized, amplified, and carried forward. The 96-well format was used to process pooled phage libraries rather than to assign one candidate clone to each well; accordingly, it did not imply a 96-candidate downstream screening budget.

PacBio Revio reads were translated from the detected start methionine and parsed into paired VH and VL segments using the known scFv linker. Reads with premature stops, frameshifts, incomplete translations, or unparseable chains were removed. Designed NNK-derived TAG codons in the amber-suppressible context were translated as glutamine (Q). Exact normalized paired amino-acid keys, VH|VL, were collapsed, and duplicate counts were summed separately for R1, R2, and R3 before mutation-position signatures were generated.

#### 1.2 Split construction and candidate-manifest handling

Equal-length candidate and parental chains were compared position by position; unequal-length chains used Needleman–Wunsch global alignment with match, mismatch, and gap scores of +2, −1, and −2. Mutation-position signatures stored chain identity and one-based position only: substitutions and deletions used the parental coordinate, insertions used the candidate coordinate when no parental coordinate existed, and residues and event identities were omitted. An empty signature was `NO_MUT`. Exact serialization and sorting rules are retained in the versioned implementation.

Eligible unmeasured candidates with identical signatures remained in the same target-specific `GroupShuffleSplit` partition. With validation fraction 0.1 and split seed 42, the fixed partitions contained 294,595 fit and 32,654 validation rows for FAF1 and 133,265 fit and 14,804 validation rows for VEGFR. Audits found zero mutation-position-signature overlap between fit and validation; memberships were fixed across methods and stochastic seeds.

Candidate manifests were defined from total count before evaluation and retained all cutoff-count ties. The primary top-5% manifests were fixed independently of apparent-affinity values and recovery outcomes and contained 16,323 FAF1 and 7,487 VEGFR candidates, with minimum total counts of 6 and 18. Reference top-1% manifests contained 3,509 and 1,481 candidates, whereas top-10% manifests contained 37,024 and 16,309; the FAF1 top-1% manifest contained only three of seven evaluation-panel clones. The primary selection depth was fixed before recovery evaluation at 384 candidate selections as an operational high-throughput downstream-screening benchmark motivated by 384-well assay formats. This budget did not represent the historical number of colonies screened, the 96-well pooled-panning format, or an assumption that 384 unique candidates would occupy one physical plate after controls and replicates. AbPACER produces a complete ranking that can be truncated to a different laboratory-specific budget; the primary claims of this study are specific to the reported 384-candidate endpoint. Budgets of 1,000, 2,000, and 5,000 were retained as secondary hypothetical scaling analyses.

NGS-matched quantitatively measured clone keys were excluded from split construction, fitting, internal validation, neighbor-reference construction, and checkpoint selection. Their paired sequences and counts remained in the complete collapsed pools, and any such clone present in a frozen candidate manifest was scored and ranked with all other candidates. Apparent-affinity values did not enter fitting, checkpoint selection, scoring, or within-method ordering. Reference and control entries were not counted toward recovery. Exact manifest and retrospective audit results are reported in Sections 2 and 4.

#### 1.3 Count and trajectory evidence

For clone $i$ in round $r$, with count $c_{r,i}$ and round total $C_r = \sum_i c_{r,i}$, $N$ was the number of exact paired VH|VL clone keys in the target-specific collapsed union of the R1–R3 outputs. A clone not observed in a given round was assigned $c_{r,i} = 0$ before application of the fixed pseudocount $p = 1$. The smoothed frequency was

$$\pi_{r,i} = \frac{c_{r,i} + p}{C_r + pN}. \tag{1}$$

The log enrichment from earlier round $a$ to later round $b$ and its sampling-noise-normalized form were

$$e_{b/a,i} = \log \pi_{b,i} - \log \pi_{a,i}, \qquad Z_{b/a,i} = \frac{e_{b/a,i}}{\sqrt{1/(c_{b,i} + p) + 1/(c_{a,i} + p)}}. \tag{2}$$

Natural logarithms were used throughout the enrichment calculation. The peak statistic was $T_{\text{peak},i} = \max(Z_{2/1,i}, Z_{3/1,i})$; $Z_{3/2,i}$ was retained separately. Total count $C_i$ was the cumulative R1–R3 support and primary non-learned baseline, not an affinity label. After training-set standardization, total count and peak trajectory were combined with fixed equal coefficients to define $O_i$.

$$O_i = \frac{1}{2} z_{\text{train}}[\log(1 + C_i)] + \frac{1}{2} z_{\text{train}}[T_{\text{peak},i}] .$$

The 1:1 weighting was fixed across targets, methods, and seeds and was not tuned against retrospective recovery; it did not imply biological equivalence, independence, or optimal weighting. Thus, $O_i$ was the affinity-label-blind fitting target and source of neighbor evidence rather than an apparent-affinity estimate.

Round-specific counts, $Z_{2/1,i}$, $Z_{3/1,i}$, $Z_{3/2,i}$, peak trajectory, and total count were deterministic controls, with higher values indicating higher priority. All deterministic and learned rankings used the frozen paired VH|VL key to resolve exact score ties.

#### 1.4 Parent-aware representation and neighbor-aware base scoring

Each explicit parent-to-candidate mutation token encoded chain, parent-referenced framework/CDR region, one-based model position, parent and candidate residues, and standardized differences in hydropathy, charge, molecular mass, polar-or-charged status, and aromaticity; a no-mutation token handled unchanged candidates. Frozen IgBERT encoded parent and candidate VH [SEP] VL sequences separately. Candidate and parent residue embeddings and their signed and absolute differences were projected and merged with the explicit token, while a global token combined candidate and parent mean embeddings and their differences. The mutation encoder, context projections, merge network, and transformer were trainable, and the transformer produced query vector $q_i$. Architecture dimensions and fixed fitting constants are summarized in Supplementary Table S1, Panel B.

Base neighbors were selected by cosine similarity in normalized signed global parent-to-candidate IgBERT-difference space. During holdout fitting, training queries used the fixed training rows as the reference pool with self-matches removed, whereas validation queries used training rows only. Quantitatively measured clone keys were never eligible as references. This attention retrieval was distinct from the residual local-family graph. Attention inputs were the neighbor difference vector, cosine similarity and distance, Gaussian weight, and $O_j$; a learned null-neighbor option allowed the model to reject an uninformative retrieval set. Four auxiliary values summarized attention-weighted $O_j$, null weight, effective neighbor count, and a similarity–distance contrast. With these values denoted by $a_i$, the base feature was

$$h_i = [q_i, n_i, a_i].$$

**Global base pairwise loss.** For a sampled mini-batch, define

$$\Delta O_{ij} = O_i - O_j, \qquad \mathcal{V}_G = \{(i,j) : |\Delta O_{ij}| > 10^{-8}\}.$$

For each retained pair,

$$r^G_{ij} = |\Delta O_{ij}| \left\{\max(O_i, O_j, 0) + 10^{-3}\right\},$$

$$Z_G = \max\left\{\text{stopgrad}\left(\text{mean}_{\mathcal{V}_G} r^G\right), 10^{-6}\right\}, \qquad w^G_{ij} = \frac{r^G_{ij}}{Z_G},$$

and

$$\mathcal{L}_G = \text{mean}_{\mathcal{V}_G}\, w^G_{ij}\, \text{softplus}\left[-\operatorname{sign}(\Delta O_{ij})(b_i - b_j)\right].$$

Pairs outside $\mathcal{V}_G$ were omitted, the normalization was detached from gradient computation, and there was no pointwise regression term. Sampling probabilities were separate from the multiplicative weight $w^G_{ij}$. Base checkpoints were selected by validation Spearman correlation with $O_i$. During holdout fitting, the standardization moments used to construct $O_i$ and $O_j$ were estimated from the fixed unmeasured training rows and applied unchanged to validation rows. A newly initialized final Base model was then refit on all eligible unmeasured rows for the selected number of epochs. For this final refit and final candidate scoring, the standardization moments and reference graph were rebuilt from all eligible unmeasured rows, including the former validation rows; each unmeasured query was removed from its own reference set. Frozen IgBERT remained unchanged, and each stage-specific neighbor graph was fixed during its corresponding fit. Predefined numerical-health checks rejected nonfinite values, near-constant scores, or all-null attention before residual fitting.

Resolved optimization, pair-sampling, checkpoint, and final-refit constants are summarized in Supplementary Table S1, Panels C and D.

#### 1.5 Local-family graph and bounded residual fitting

**Frozen-IgBERT parent-delta relation.** Let $\bar{\mathbf{h}}_i$ be the mean final-layer frozen-IgBERT state over VH and VL residue tokens, excluding special tokens, and define parental $\bar{\mathbf{h}}_0$ identically. The normalized parent-relative vector and cosine distance were

$$\mathbf{u}_i = \frac{\bar{\mathbf{h}}_i - \bar{\mathbf{h}}_0}{\|\bar{\mathbf{h}}_i - \bar{\mathbf{h}}_0\|_2}, \qquad d_{ij} = 1 - \mathbf{u}_i^{\mathsf{T}} \mathbf{u}_j.$$

The Gaussian relation was

$$k^{\text{IgBERT}}_{ij} = \text{safeexp}\left[-\frac{1}{2}\left(\frac{d_{ij}}{\sigma_t}\right)^2\right],$$

where target-specific $\sigma_t$ values are listed in Supplementary Table S1, Panel B. Here safeexp denotes the numerically clipped exponential specified in the versioned implementation. The squared quantity was cosine distance itself; no Gaussian density prefactor or row normalization was applied.

**Continuous count similarity.** With $C_i = c_{R1,i} + c_{R2,i} + c_{R3,i}$,

$$k^{\text{count}}_{ij} = \text{safeexp}\left[-\frac{|\log(1 + C_i) - \log(1 + C_j)|}{0.45}\right].$$

This was an absolute-difference exponential kernel.

**Exact-mutation overlap.** Let $M_i$ be the stored mutation-tuple set

$$(\text{chain}, \text{one-based model coordinate}, \text{parental residue}, \text{candidate residue}).$$

The coordinate was parental when available and candidate-referenced for an insertion; insertions and deletions used `-` for the absent residue, and `NO_MUT` was omitted. Define

$$m_{ij} = |M_i \cap M_j|,$$

$$J_{ij} = \begin{cases} \dfrac{m_{ij}}{|M_i \cup M_j|}, & |M_i \cup M_j| > 0, \\ 0, & |M_i \cup M_j| = 0, \end{cases} \qquad A_{ij} = \frac{m_{ij}}{\max(|M_i|, 1)},$$

$$k^{\text{mutation}}_{ij} = 1 + 2\max(J_{ij}, A_{ij}).$$

Because $A_{ij}$ measured anchor coverage, this relation need not be symmetric.

**Combined local relation and winsorized support.** The unnormalized relation product was

$$w_{ij} = k_{ij}^{\mathrm{IgBERT}} k_{ij}^{\mathrm{count}} k_{ij}^{\mathrm{mutation}}.$$

No row normalization was applied; nonfinite or nonpositive relations were discarded after self-removal.

For anchor $i$, let $\mathcal{I}_i$ contain the positive finite relations in its self-excluded local graph and let

$$\mathcal{N}_i = \{j \in \mathcal{I}_i : O_j \text{ is finite}\}.$$

For nonempty $\mathcal{N}_i$, the unweighted neighbor-target quantiles were

$$Q_i^- = Q_{0.05}\{O_j : j \in \mathcal{N}_i\}, \qquad Q_i^+ = Q_{0.95}\{O_j : j \in \mathcal{N}_i\},$$

$$\widetilde{O}_{j|i} = \mathrm{clip}(O_j, Q_i^-, Q_i^+), \qquad L_i = \frac{\sum_{j \in \mathcal{N}_i} w_{ij} \widetilde{O}_{j|i}}{\sum_{j \in \mathcal{N}_i} w_{ij}}.$$

The 5th and 95th percentiles were unweighted. Nonfinite targets were omitted, and an empty effective neighborhood used the implemented fallback $L_i = O_i$.

**Bounded residual and local anchor-pair loss.** The frozen base score was refined by

$$\delta_i = \Delta_{\max} \tanh(g_\theta(h_i)), \qquad s_i = b_i + \delta_i, \qquad \Delta_{\max} = 1.0.$$

The score $s_i$ was the final AbPACER ranking score.

The residual head was initialized to zero and the base parameters remained frozen during residual fitting. For anchor mini-lists, define

$$\mathcal{V}_L = \{(i, j) : j \text{ is a masked-in neighbor of anchor } i, w_{ij} > 0\}.$$

The raw and normalized local pair weights were

$$a_{ij} = w_{ij}^{0.5} \max(|L_i - L_j|, 10^{-6}),$$

$$Z_L = \max\left\{\mathrm{stopgrad}\left(\mathrm{mean}_{\mathcal{V}_L} a\right), 10^{-6}\right\}, \qquad \widehat{a}_{ij} = \frac{a_{ij}}{Z_L},$$

and the temperature-1.0 training loss was

$$\mathcal{L}_{\mathrm{local}} = \mathrm{mean}_{\mathcal{V}_L} \widehat{a}_{ij} \, \mathrm{softplus}\left[-\mathrm{sign}(L_i - L_j)(s_i - s_j)\right].$$

There was no positive minimum target-gap filter, and exact target ties produced zero score gradient. Residual checkpoints used held-out unweighted anchor-pair softplus loss; epoch 0, corresponding to no residual correction, was eligible. If no trained checkpoint improved on epoch 0, the Base score was retained. All six primary phage-display runs selected an improved nonzero checkpoint. A newly initialized residual head was then refit on all eligible unmeasured local-family rows for the selected number of epochs while the Base remained frozen.

**Local-region and list construction.** The local-family region was calculated before training exclusion on the full deduplicated pool using descending average ranks:

$$100 \, \frac{\mathrm{rank}_{\mathrm{average,descending}}(C_i)}{N} \leq 5.$$

Rows excluded under Section 1.2 were removed afterward. Pre-exclusion memberships matched the primary manifests: 16,323 FAF1 keys with $C_i \geq 6$ and 7,487 VEGFR keys with $C_i \geq 18$, including cutoff ties. Removing the parent and matched measured clones left 16,315 and 7,483 active rows.

Local graphs were distinct from base attention. During holdout fitting, training anchors retrieved self-excluded training-local candidates, whereas validation queries used training-local references only. For the final residual refit, the graph was rebuilt over all eligible unmeasured local-family rows, with each row removed from its own reference set. Support used positive finite combined relations. Retrieval and mini-list caps are in Supplementary Table S1, Panel B. Deterministic sorting, numerical clipping, dtype transitions, and edge-case fallback rules are specified in the versioned implementation and resolved configurations.

### Supplementary Table S1. Signal glossary and fixed architecture and fitting constants

#### Panel A. Signal and notation glossary

| Symbol | Meaning | Role in the workflow |
|---|---|---|
| $C_i$ | Total R1–R3 read count | Cumulative empirical read support, candidate-manifest construction variable, and primary non-learned baseline |
| $T_{\text{peak},i}$ | Strongest noise-normalized R1-to-later-round enrichment | Selection-trajectory signal |
| $O_i$ | Equal-weight standardized combination of count and peak trajectory | Affinity-label-blind global target and neighbor evidence |
| $q_i$ | Transformer summary of the parent-relative candidate sequence | Query-clone representation |
| $n_i$ | Attention-weighted representation of retrieved neighbors | Neighbor context |
| $a_i$ | Four scalar neighborhood summaries | Auxiliary evidence supplied to scoring heads |
| $b_i$ | Stage-1 scalar output | Base ranking score |
| $L_i$ | Winsorized, relation-weighted neighbor $O_j$ support | Local neighbor support |
| $\delta_i$ | Bounded stage-2 correction | Local adjustment to the base score |
| $s_i$ | $b_i + \delta_i$ | Final AbPACER ranking score |

None of $O_i$, $b_i$, $L_i$, $\delta_i$, or $s_i$ is a measured or calibrated affinity value. $O_i$ is the global affinity-label-blind fitting target and source of neighbor evidence, whereas $s_i$ is the final learned score used to rank phage-display candidates directly.

#### Panel B. Architecture and fitting constants

| Component | Fixed value | Scope |
|---|---|---|
| Explicit mutation representation | 128 dimensions | Per parent-to-candidate mutation token |
| IgBERT residue-context projection | 64 dimensions | Candidate, parent, signed-difference, and absolute-difference projection |
| Context-aware mutation token | 128 dimensions | Merged explicit and IgBERT residue context |
| Global-context token | 128 dimensions | Candidate/parent mean context and differences |
| Transformer encoder | 2 layers; 4 heads; 128-dimensional token width | Global and mutation-token encoder producing $q_i$ |
| Base-neighbor retrieval cap | 32 | Attention-stage unmeasured references |
| Local-neighbor retrieval cap | 64 | Self-excluded local-family candidates |
| Residual mini-list size | Anchor plus at most 11 neighbors; at least 3 total members | Local residual fitting |
| IgBERT relation bandwidth $\sigma_t$ | FAF1: 0.007346392376382136; VEGFR: 0.002494733897047373 | Target-specific local relation |
| Count-kernel scale | 0.45 | Absolute log-count-difference kernel |
| Winsorization quantiles | 0.05 and 0.95 | Unweighted local $O_j$ support |
| Residual bound $\Delta_{\max}$ | 1.0 | Bounded correction $\delta_i$ |
| Local-loss temperature | 1.0 | Anchor-pair softplus loss |
| Phage maximum training epochs | Base: 20; residual: 15 | Phage holdout checkpoint selection before final refit |
| AlphaSeq fixed fitting protocol | Pointwise mean-squared error; global validation Pearson checkpoint selection | One pooled model per seed; library-specific validation-Spearman bandwidth selection |

Base and residual fitting used FP32. Numerically sensitive relation and normalization calculations used higher precision before conversion to the model input dtype. Exact dtype transitions and numerical safeguards are documented in the versioned implementation.

#### Panel C. Resolved optimization constants

| Benchmark / method | Stage | Optimizer | Learning rate | Weight decay | Batch unit and size | Dropout / gradient clipping | Epoch and checkpoint rule |
|---|---|---|---|---|---|---|---|
| Phage / AbPACER | Base | AdamW | $1{\times}10^{-3}$ | $1{\times}10^{-4}$ | 64 sampled pairs | 0.10 / global norm 5.0 | Maximum 20; validation Spearman; patience 3; minimum delta $10^{-4}$; exact seed-wise selected/executed epochs in note C1. |
| Phage / AbPACER | Residual | AdamW, residual head only | $3{\times}10^{-4}$ | $1{\times}10^{-4}$ | 16 anchor mini-lists; 188 steps/epoch | Head 0; frozen Base in evaluation mode / 5.0 | Maximum 15 trained epochs; minimum held-out anchor-pair loss with epoch 0 eligible; patience 3; minimum decrease $10^{-4}$; note C1. |
| Phage / Ens-Grad CNN | Six constituents, separately optimized | AdamW | $1{\times}10^{-3}$ | $1{\times}10^{-4}$ | 64 sampled pairs | 0.30 in five members; 0.35 in `cnn64x2_k3` / 5.0 | Maximum 20 per member; validation Spearman; patience 3; minimum delta $10^{-4}$; exact member-wise epochs in note C2. |
| Phage / A2Binder-HL | Full H/L model | AdamW, two groups | Encoders $2{\times}10^{-5}$; head/CNN $1{\times}10^{-4}$ | $1{\times}10^{-4}$ | 4 physical / 64 effective pairs; accumulation 16 | H/L hidden and attention 0.10; custom head 0 / 5.0 | Maximum 20; validation Spearman; patience 3; minimum delta $10^{-4}$; note C1. |
| Phage / AbAffinity | Full ESM-2 and linear head | AdamW | $1{\times}10^{-5}$ | 0 | 4 physical / 64 effective pairs; accumulation 16 | No stochastic dropout / 5.0 | Maximum 20; validation Spearman; patience 3; minimum delta $10^{-4}$; note C1. |
| AlphaSeq / AbPACER-MSE | Base | AdamW | $2{\times}10^{-4}$ | $1{\times}10^{-4}$ | 128 clone rows | 0.10 / global norm 5.0 | Maximum 100; pooled validation Pearson; patience 10; improvement $> 10^{-12}$; selected/executed: seed 42, 15/25; 43, 16/26; 44, 17/27. |
| AlphaSeq / AbPACER-MSE | Residual | Fresh AdamW; selected Base frozen | $2{\times}10^{-4}$ | $1{\times}10^{-4}$ | 128 clone rows | Head 0; frozen Base 0.10 / 5.0 | Maximum 100; pooled validation Pearson; patience 10; improvement $> 10^{-12}$; selected/executed: seed 42, 5/15; 43, 3/13; 44, 6/16. |
| AlphaSeq / Ens-Grad CNN | Six constituents, separately optimized | AdamW | $1{\times}10^{-3}$ | $1{\times}10^{-4}$ | 512 pooled clone rows | 0.30 in five members; 0.35 in `cnn64x2_k3` / 1.0 | Maximum/executed 40/40 for all 18 fits; strict validation-Pearson maximum; minimum 5 epochs and patience 8; exact selected epochs in note C3. |
| AlphaSeq / A2Binder | Seed-specific continuation | AdamW, two groups | Encoders $2{\times}10^{-5}$; heads $1{\times}10^{-4}$ | $1{\times}10^{-4}$ | 16 rows × accumulation 8 = 128 effective | H/L hidden and attention 0.10; antigen 0; heads 0 / 1.0 | Maximum/executed 12/12; no early stopping; strict validation-Pearson maximum; selected epoch 2 in every seed. |
| AlphaSeq / AbAffinity | Full ESM-2 and linear predictor | Adam | $1{\times}10^{-5}$ | 0 | 32 rows × accumulation 4 = 128 effective | No stochastic dropout / 1.0 | Maximum/executed 100/100; no early stopping; strict validation-Pearson maximum; selected epochs 11, 11, and 10 for seeds 42, 43, and 44. |

C1: Values are selected/executed epochs for seeds 42, 43, and 44, respectively. Phage AbPACER Base: FAF1 10/13, 8/11, 12/15; VEGFR 7/10, 1/4, 1/4. Residual: FAF1 8/11 for every seed; VEGFR 6/9 for every seed. A2Binder-HL: FAF1 10/13, 11/14, 6/9; VEGFR 9/12, 6/9, 5/8. AbAffinity: FAF1 10/13, 9/12, 10/13; VEGFR 8/11, 8/11, 6/9.

C2: Phage Ens-Grad architecture order was `cnn32_k5`, `cnn64_k5`, `cnn32x2_k5`, `cnn32_k3`, `cnn64x2_k3`, and `cnn_emb_32x1_16`. Selected epochs were FAF1 seed 42 [20,20,12,20,1,4], seed 43 [9,20,3,15,20,2], seed 44 [20,16,4,20,2,2]; VEGFR seed 42 [15,20,10,20,20,20], seed 43 [20,14,20,20,20,19], seed 44 [20,17,19,20,20,3]. Executed member horizons are retained in the machine-readable trace.

C3: In the same architecture order, AlphaSeq Ens-Grad selected epochs were seed 42 [39,34,39,40,36,39], seed 43 [39,40,40,40,37,40], and seed 44 [39,38,39,35,40,40].

All reported training used no learning-rate scheduler or warmup. Phage AbPACER used FP32. Phage comparators used BF16 autocast with FP32 reductions, except the predefined same-seed FP32 retry for VEGFR seed 44 Ens-Grad member `cnn64_k5`. AlphaSeq AbPACER-MSE used FP16 autocast with GradScaler and FP32 parameters/optimizer; AlphaSeq Ens-Grad used FP32; AlphaSeq A2Binder and AbAffinity used BF16 autocast with FP32 parameters/optimizer. Phage global stages used the documented weighted pairwise softplus loss, the phage residual used the local anchor-pair softplus loss at temperature 1.0, and AlphaSeq stages used pointwise MSE. Where an Adam/AdamW call did not override them, betas $(0.9, 0.999)$, epsilon $10^{-8}$, and AMSGrad false were verified against the retained PyTorch environment (2.12.0, or 2.8.0 for AlphaSeq A2Binder).

**Panel D. Resolved sampling and execution protocol**

| Benchmark / method | Sampling unit | Per-batch or per-epoch schedule | Validation construction | Early stopping / patience | Final-refit policy |
|---|---|---|---|---|---|
| Phage / AbPACER Base | Presampled ordered anchor–opponent pairs | 64 pairs/step; 16,000 pairs and 250 steps/full epoch; same serialized schedules as all global-pairwise comparators | All fixed validation rows scored each epoch: FAF1 32,654 and VEGFR 14,804; Spearman with $O_i$ | Patience 3; minimum delta $10^{-4}$ | Fresh Base and AdamW at seed +100000; all eligible unmeasured rows; exactly the selected epoch count |
| Phage / AbPACER residual | Anchor-centered mini-lists | 16 lists/step; 188 steps and 3,008 list draws/full epoch; anchor plus at most 11 relation-ranked neighbors, minimum 3 total | Fixed lists: FAF1 1,549 lists/16,776 realized anchor pairs; VEGFR 725/7,955; epoch-0 loss included | Patience 3 trained epochs; minimum loss decrease $10^{-4}$ | Fresh zero-initialized head and AdamW at seed +300000; final Base frozen; exactly the selected nonzero epoch count |
| Phage / Ens-Grad CNN | Shared presampled global pairs; six model members | 64 pairs/step; 16,000/full epoch per member; no accumulation | All fixed validation rows; Spearman with $O_i$, separately per member | Patience 3; minimum delta $10^{-4}$ | Fresh model and AdamW per member at member seed +100000; all eligible unmeasured rows; member-specific selected epoch count |
| Phage / A2Binder-HL | Shared presampled global pairs | 4 physical pairs/microbatch; accumulation 16; 64 effective; 16,000/full epoch | All fixed validation rows; Spearman with $O_i$ | Patience 3; minimum delta $10^{-4}$ | Fresh full model and AdamW at seed +100000; all eligible unmeasured rows; selected epoch count |
| Phage / AbAffinity | Shared presampled global pairs | 4 physical pairs/microbatch; accumulation 16; 64 effective; 16,000/full epoch | All fixed validation rows; Spearman with $O_i$ | Patience 3; minimum delta $10^{-4}$ | Fresh full model and AdamW at seed +100000; all eligible unmeasured rows; selected epoch count |
| AlphaSeq / AbPACER-MSE Base | Clone-row batches in library-tagged chunks | Shuffle rows within each library, form 128-row chunks, then shuffle tagged chunks; final partial chunks retained | Global Pearson over all 9,025 fixed validation rows; fit rows are the only neighbor references; within-library bandwidth selection uses up to 4,000 validation queries | Patience 10; improvement $> 10^{-12}$ | None: selected Base is frozen for residual fitting; no fit-plus-validation refit |
| AlphaSeq / AbPACER-MSE residual | Same pooled clone-row chunk schedule | 128 rows/step; residual row-order RNG uses seed +11; no pair, anchor, or mini-list sampling | Global Pearson over the same 9,025 validation rows | Patience 10; improvement $> 10^{-12}$ | None: selected residual state directly generated frozen predictions |
| AlphaSeq / Ens-Grad CNN | Shuffled pooled clone rows, separately per member | 512 rows/batch; six architecture-defined constituents; all ran 40 epochs | Global Pearson over all 9,025 validation rows, separately per member | Configured minimum 5 and patience 8; all fits completed 40 epochs | None: six selected-checkpoint predictions were averaged |
| AlphaSeq / A2Binder | Shuffled pooled clone rows; four deterministically seeded workers | 16 rows/microbatch; accumulation 8; 128 effective; 12 epochs | Global Pearson over all 9,025 validation rows | No early stopping or minimum-delta offset | No final refit: all seeds loaded one common seed-42-trained precursor, then used a fresh AdamW and seed-specific 12-epoch continuation; selected continuation checkpoint predicted directly |
| AlphaSeq / AbAffinity | Shuffled pooled clone rows; four seeded workers | 32 rows/microbatch; accumulation 4; 128 effective; random worker-seeded crop only for training sequences longer than 256 residues | Global Pearson over all 9,025 validation rows; validation/test used center crop | No early stopping or minimum-delta offset | None: selected checkpoint reloaded and predicted directly |

Global phage-pair schedules were presampled separately for holdout and final refit. Anchor probability was proportional to $1 + \max(O_i, 0)/\operatorname{mean}\{\max(O, 0)\}$. For each anchor, 96 opponent candidates were drawn with replacement; the implemented choice weight combined count ambiguity, target gap, and top-target support. Self-pairs were excluded, non-ties were preferred when available, anchor/opponent orientation was retained, and repeated schedule pairs were allowed. Holdout schedule seeds were stochastic seed +17; final-refit schedule seeds were stochastic seed +100017. The complete formula and code-line trace are provided in the machine-readable audit.

AlphaSeq A2Binder provenance requires care: the common precursor was configured for 15 epochs but retains seven trace rows and an epoch-3 best state; its termination reason is not recoverable. Each reported seed then completed the resolved 12-epoch continuation from that common state with a fresh optimizer. Random seed and worker policies are resolved for all AlphaSeq methods, but explicit deterministic-kernel enforcement is not recoverable from retained run records. Four-library-specific AbPACER-MSE used the pooled constants except for four independently initialized library models, per-library fit normalization, no inter-library chunk mixing, and per-library validation-Pearson checkpointing.

##### 1.6 AlphaSeq target transformation and neighbor-selection protocol

These procedures applied only to AlphaSeq and used no phage-display counts, trajectories, $O_i$, or phage neighbor evidence. Every method used the immutable 51,139/9,025/11,670 fit/validation/test assignments, seeds 42–44, squared-error training, validation-Pearson checkpoint selection, raw test predictions, and no calibration. Test labels were not used for bandwidth selection, fitting, checkpoint selection, or calibration.

AbPACER-MSE, AbAffinity, and A2Binder used one pooled model per seed; Ens-Grad CNN averaged six pooled constituent regressors. AbPACER-MSE shared one parameter set across libraries while using the appropriate parental clone, parent-relative features, frozen IgBERT context, and within-library feature and neighbor providers, without a library-identity embedding.

Bandwidth was selected separately within each library by maximizing Spearman correlation with the validation $K_D$-derived target. Validation rows were queries and fit rows were the only neighbor references; validation and test rows were never neighbor references. Test targets remained inaccessible until evaluation of frozen predictions.

For AlphaSeq, let $y_i = \log_{10}(K_{D,i})$. The mean $\mu_{\text{fit}}$ and standard deviation $\sigma_{\text{fit}}$ were calculated only from the complete 51,139-row fit cohort, and the normalized target was

$$t_i = -\frac{y_i - \mu_{\text{fit}}}{\sigma_{\text{fit}}}.$$

Fit-derived statistics were applied unchanged to validation and test rows. Validation targets supported bandwidth and checkpoint selection but not model fitting. The shared Base model and bounded residual head were trained sequentially by pointwise mean-squared error, with Base parameters frozen during residual fitting. Predictions were returned to the original affinity scale as

$$\widehat{y_i} = \mu_{\text{fit}} - \sigma_{\text{fit}}\widehat{t_i}.$$

Checkpoint selection used global validation Pearson over all 9,025 validation rows. AbAffinity fine-tuned ESM-2 t33 650M end to end, A2Binder used heavy-, light-, and antigen-sequence branches with the fixed HR2 peptide, and Ens-Grad CNN used six predefined one-hot Conv1D regressors.

The primary endpoint ranked all 11,670 fixed-test variants and measured overlap between each predicted top 384 and the 384 variants with lowest measured $K_D$; precision@384 and recall@384 were identical. Secondary analyses recovered the true top-1% (117), top-5% (584), and top-10% (1,167) sets among the same 384 predictions. The selection depth matched the primary 384-candidate fixed-budget benchmark used in the phage-display arm, while supervision and denominators remained distinct. Exact ties used the immutable test-row key. Section 7 reports the detailed results. Within-library analyses reused frozen predictions without retraining, checkpoint reselection, or recalibration.

### 2 Full-pool statistics, frozen candidate manifests, and retrospective measurement audit

This section describes the complete sequencing pools, frozen candidate manifests, and retrospective evaluation panels.

Main Table 1 provides a condensed summary of full-pool scale, singleton burden, and the primary candidate manifests. Supplementary Table S2 extends this summary with cumulative low-count statistics and the top-1%, top-5%, and top-10% manifest definitions used to document candidate-universe scale and panel coverage around the primary choice.

The top-5% manifests defined the primary candidate universes, and the primary selection depth of 384 served as the practical fixed-budget benchmark. The top-1% and top-10% manifests are reported as narrower and broader reference universes rather than as alternative primary analyses.

#### Supplementary Table S2. Extended full-pool sparsity statistics and frozen candidate manifests

##### Panel A. Full-pool phage-display statistics

| Target | Unique clones | R1 reads | R2 reads | R3 reads | Total reads | Count = 1 | Count ≤ 2 | Count ≤ 5 |
|---|---|---|---|---|---|---|---|---|
| FAF1 | 327 258 | 128 445 | 472 725 | 117 998 | 719 168 | 206 584 (63.1%) | 265 092 (81.0%) | 310 935 (95.0%) |
| VEGFR | 148 073 | 216 181 | 489 642 | 449 405 | 1 155 228 | 79 708 (53.8%) | 104 455 (70.5%) | 126 541 (85.5%) |

##### Panel B. Frozen candidate manifests

| Target | Candidate universe | Candidates | Primary panel members | Analysis role |
|---|---|---|---|---|
| FAF1 | Top 1% by total count | 3 509 | 3 | Narrow reference universe |
| FAF1 | Top 5% frozen manifest | 16 323 | 7 | Primary analysis |
| FAF1 | Top 10% frozen manifest | 37 024 | 7 | Broad reference universe |
| VEGFR | Top 1% by total count | 1 481 | 3 | Narrow reference universe |
| VEGFR | Top 5% frozen manifest | 7 487 | 3 | Primary analysis |
| VEGFR | Top 10% frozen manifest | 16 309 | 3 | Broad reference universe |

Candidate universes were defined by total count and instantiated as frozen clone-key manifests before comparative evaluation; all cutoff-count ties were retained. The top-1% FAF1 reference universe contains three members of the fixed seven-clone primary panel and therefore does not preserve the complete evaluation panel. The top-10% FAF1 and VEGFR reference universes contain approximately 2.27-fold and 2.18-fold more candidates than their primary top-5% manifests. The primary manifests remove the extensive low-count tail while retaining 16 323 FAF1 and 7 487 VEGFR candidates, all members of the fixed primary panels, and candidate spaces substantially larger than the 384-candidate assay budget. Exact clone keys and manifest hashes will be included in the planned archival release.

Supplementary Table S3 reports all quantitative scFv–Fc concentration–response measurements available from the historical workflow. Only clones that advanced from the preceding scFv–Fc screening step underwent quantitative scFv–Fc measurement, and no quantitatively measured clone was excluded on the basis of its binding result. All clones in this historical quantitative set had lower target-specific ELISA-derived apparent affinity values than their respective parental clones.

The table contains the complete available quantitative measurement set. Supplementary Figure S1 shows the subset for which historical curve-level fit outputs were available.

#### Supplementary Table S3. Retrospective measurement metadata and evaluation membership

| Target | Clone | Parental clone | Parental apparent affinity | Clone apparent affinity | Parental/clone ratio | Additional apparent affinity | Total count | NGS match | Top-5% | Primary panel |
|---|---|---|---|---|---|---|---|---|---|---|
| FAF1 | FAF1_AM_#18 | FAF1_#1 | 43.1 pM | 21.6 pM | 1.99 | – | – | no | no | no |
| FAF1 | FAF1_AM_#8 | FAF1_#1 | 43.1 pM | 21.8 pM | 1.98 | – | 7 | yes | yes | yes |
| FAF1 | FAF1_AM_#20 | FAF1_#1 | 43.1 pM | 27.7 pM | 1.56 | – | 891 | yes | yes | yes |
| FAF1 | FAF1_AM_#10 | FAF1_#1 | 43.1 pM | 27.3 pM | 1.58 | – | 38 | yes | yes | yes |
| FAF1 | FAF1_AM_#6 | FAF1_#1 | 43.1 pM | 30.2 pM | 1.43 | – | 4 | yes | no | no |
| FAF1 | FAF1_AM_#12 | FAF1_#1 | 43.1 pM | 35.3 pM | 1.22 | – | 6 | yes | yes | yes |
| FAF1 | FAF1_#134 | FAF1_#1 | 43.1 pM | 30.3 pM | 1.42 | – | 22 | yes | yes | yes |
| FAF1 | FAF1_#138 | FAF1_#1 | 43.1 pM | 27.8 pM | 1.55 | – | 6 | yes | yes | yes |
| FAF1 | FAF1_#228 | FAF1_#1 | 43.1 pM | 27 pM | 1.60 | – | 9 | yes | yes | yes |
| VEGFR | AM-#3 | #163 | 20.7 nM | 3.35 nM | 6.18 | VEGFR-1: 2.59 nM | 6 638 | yes | yes | yes |
| VEGFR | AM-#63 | #163 | 20.7 nM | 1.77 nM | 11.69 | VEGFR-1: 1.22 nM | 868 | yes | yes | yes |
| VEGFR | AM-#83 | #163 | 20.7 nM | 6.54 nM | 3.17 | VEGFR-1: 1.82 nM | 371 | yes | yes | yes |

The parental/clone ratio was calculated from the underlying fit values as parental apparent affinity divided by clone apparent affinity; values greater than 1 indicate a lower apparent-affinity value for the clone. Because the displayed midpoint values are rounded, a ratio may differ slightly from a value recomputed directly from the displayed midpoint. The VEGFR parental clone #163 had an additional VEGFR-1 apparent affinity value of 2.58 nM; VEGFR-1 clone apparent affinity values are descriptive and were not used to define target-specific recovery. FAF1_AM_#18 did not match the final collapsed paired VH|VL NGS key. FAF1_AM_#6 matched the full pool but fell outside the primary top-5% manifest. Measured clones were not treated as negative examples, and apparent-affinity values did not enter fitting, checkpoint selection, score computation, or within-method candidate ordering.

**A** **FAF1: parent #1 and representative affinity-matured clones**

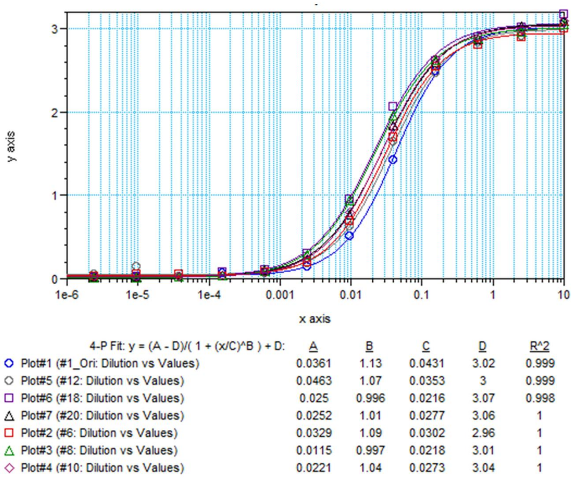

| Clone | Apparent midpoint (nM) | Parent/clone ratio |
|---|---|---|
| Parent #1 | 0.0431 | - |
| AM-#12 | 0.0353 | 1.22 |
| AM-#6 | 0.0302 | 1.43 |
| AM-#10 | 0.0273 | 1.58 |
| AM-#20 | 0.0277 | 1.56 |
| AM-#8 | 0.0218 | 1.98 |
| AM-#18 | 0.0216 | 1.99 |

**B** **VEGFR-2: parent #163 and affinity-matured clones AM-#3, AM-#63, and AM-#83**

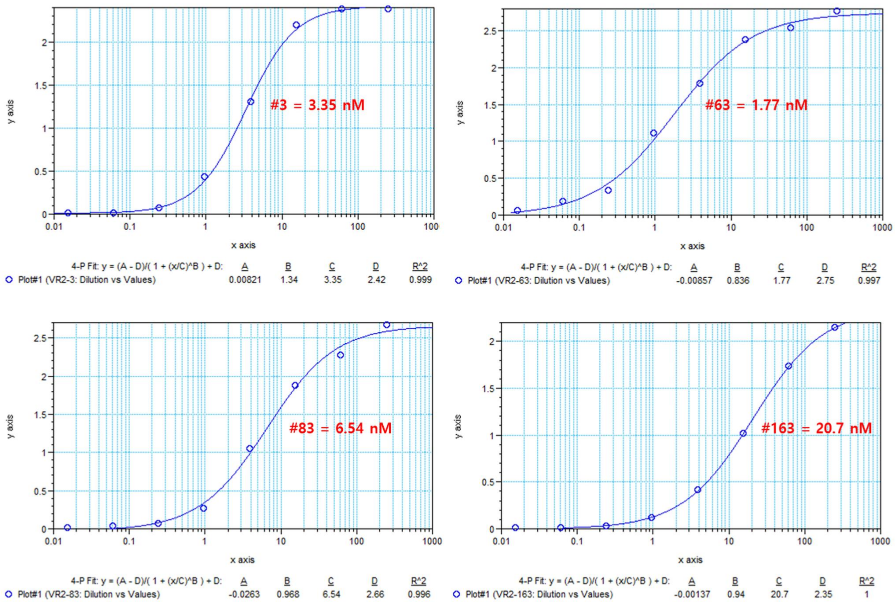

**Figure S1:** Concentration-dependent scFv–Fc binding ELISA curves underlying the retrospective evaluation panel. (A) FAF1 parental clone #1 and affinity-matured clones for which historical curve-level fit outputs were available. The displayed set includes four clones in the fixed primary FAF1 evaluation panel (AM-#8, AM-#10, AM-#12, and AM-#20) and two additional quantitatively measured clones (AM-#6 and AM-#18). (B) VEGFR-2 binding curves for parental clone #163 and affinity-matured clones AM-#3, AM-#63, and AM-#83. Source concentration–response outputs and the corresponding four-parameter logistic fits are shown. Across all panels, the horizontal axis represents scFv–Fc concentration on a logarithmic scale, and the vertical axis represents the ELISA absorbance signal; the original historical output formatting and scaling were retained. The fitted midpoint was reported as an ELISA-derived apparent affinity value, with lower values indicating stronger apparent binding. FAF1 curve-level values are shown at the precision available in the historical outputs and, for the clones displayed in Panel A, are used as the authoritative apparent-affinity values in the manuscript tables. These measurements were used only for retrospective evaluation and did not enter model fitting, checkpoint selection, score computation, or within-method candidate ordering.

### 3 Parent-to-candidate mutation maps

Mutation maps are reported for the primary evaluation panels. Positions correspond to the parent-referenced model coordinates used in the analysis and should not be interpreted as curated IMGT numbering.

#### Supplementary Table S4. Parent-to-candidate mutation maps

| Target | Clone | ELISA-derived apparent affinity | Mutations from parental clone |
|---|---|---|---|
| FAF1 | FAF1_AM_#8 | 21.8 pM | VH:57:F>L; VL:38:L>M; VL:111:I>V |
| FAF1 | FAF1_AM_#20 | 27.7 pM | VH:57:F>I |
| FAF1 | FAF1_AM_#10 | 27.3 pM | VH:13:V>E; VH:57:F>L; VH:61:N>K; VH:112:Q>R |
| FAF1 | FAF1_AM_#12 | 35.3 pM | VH:56:I>V; VL:60:F>L |
| FAF1 | FAF1_#134 | 30.3 pM | VH:57:F>I; VL:35:K>M; VL:82:R>P |
| FAF1 | FAF1_#138 | 27.8 pM | VH:45:Q>H; VH:57:F>L; VL:42:L>M |
| FAF1 | FAF1_#228 | 27 pM | VH:25:K>N; VH:97:H>Y; VH:102:G>D; VL:3:V>E; VL:7:T>I; VL:8:P>Q; VL:73:G>E |
| VEGFR | AM-#3 | 3.35 nM (VEGFR-2) | VH:3:Q>G; VH:10:G>R; VH:12:E>K; VH:17:G>E; VH:56:S>I; VH:64:Q>P; VH:75:D>E; VH:104:L>M; VH:105:V>I |
| VEGFR | AM-#63 | 1.77 nM (VEGFR-2) | VH:3:Q>K; VH:12:E>K; VH:42:A>V; VH:109:N>H; VH:118:T>A; VL:43:T>P; VL:91:A>T; VL:105:T>S |
| VEGFR | AM-#83 | 6.54 nM (VEGFR-2) | VH:30:T>A; VH:95:V>M; VH:105:V>I; VL:7:P>T; VL:35:N>T; VL:50:Y>H; VL:60:P>H; VL:98:G>A; VL:101:F>V |

### 4 Deterministic controls, direct learned rankings, and clone-level audit

The four-budget analysis compares deterministic count and trajectory controls with direct frozen outputs from each learned method. All learned methods were ranked directly by model score using the same deterministic score-tie rule. No model was retrained, no checkpoint was reselected, and no calibration was fitted for this table.

#### Supplementary Table S5. Four-budget phage-display recovery

**Panel A. FAF1 evaluation-panel recovery**

| Method | 384 | 1 000 | 2 000 | 5 000 |
|---|---|---|---|---|
| R1 read count | 1/7 | 2/7 | 5/7 | 5/7 |
| R2 read count | 1/7 | 2/7 | 3/7 | 3/7 |
| R3 read count | 1/7 | 1/7 | 2/7 | 2/7 |
| Total read count | 1/7 | 2/7 | 3/7 | 3/7 |
| $Z_{2/1}$ | 1/7 | 2/7 | 2/7 | 3/7 |
| $Z_{3/1}$ | 0/7 | 0/7 | 0/7 | 0/7 |
| $Z_{3/2}$ | 0/7 | 0/7 | 0/7 | 1/7 |
| Peak trajectory | 1/7 | 2/7 | 2/7 | 2/7 |
| Direct $O_i$ ranking | 1/7 | 2/7 | 3/7 | 3/7 |
| Count–$O_i$ rank fusion | 1/7 | 2/7 | 3/7 | 3/7 |
| $O_i$ within exact-count ties | 1/7 | 2/7 | 3/7 | 3/7 |
| Ens-Grad CNN | $1.00 \pm 0.00/7$ | $2.00 \pm 0.00/7$ | $4.33 \pm 1.15/7$ | $5.00 \pm 0.00/7$ |
| A2Binder-HL | $1.67 \pm 1.15/7$ | $2.67 \pm 0.58/7$ | $3.67 \pm 0.58/7$ | $6.00 \pm 0.00/7$ |
| AbAffinity | $1.33 \pm 0.58/7$ | $2.00 \pm 1.00/7$ | $4.00 \pm 1.00/7$ | $5.67 \pm 0.58/7$ |
| AbPACER | $2.00 \pm 0.00/7$ | $2.00 \pm 0.00/7$ | $3.67 \pm 0.58/7$ | $5.33 \pm 0.58/7$ |

**Panel B. VEGFR evaluation-panel recovery**

| Method | 384 | 1 000 | 2 000 | 5 000 |
|---|---|---|---|---|
| R1 read count | 1/3 | 1/3 | 2/3 | 3/3 |
| R2 read count | 3/3 | 3/3 | 3/3 | 3/3 |
| R3 read count | 3/3 | 3/3 | 3/3 | 3/3 |
| Total read count | 3/3 | 3/3 | 3/3 | 3/3 |
| $Z_{2/1}$ | 3/3 | 3/3 | 3/3 | 3/3 |
| $Z_{3/1}$ | 3/3 | 3/3 | 3/3 | 3/3 |
| $Z_{3/2}$ | 2/3 | 3/3 | 3/3 | 3/3 |
| Peak trajectory | 3/3 | 3/3 | 3/3 | 3/3 |
| Direct $O_i$ ranking | 3/3 | 3/3 | 3/3 | 3/3 |
| Count–$O_i$ rank fusion | 3/3 | 3/3 | 3/3 | 3/3 |
| $O_i$ within exact-count ties | 3/3 | 3/3 | 3/3 | 3/3 |
| Ens-Grad CNN | $0.33 \pm 0.58/3$ | $1.00 \pm 1.00/3$ | $3.00 \pm 0.00/3$ | $3.00 \pm 0.00/3$ |
| A2Binder-HL | $1.00 \pm 0.00/3$ | $2.33 \pm 1.15/3$ | $3.00 \pm 0.00/3$ | $3.00 \pm 0.00/3$ |
| AbAffinity | $1.33 \pm 0.58/3$ | $3.00 \pm 0.00/3$ | $3.00 \pm 0.00/3$ | $3.00 \pm 0.00/3$ |
| AbPACER | $2.33 \pm 0.58/3$ | $2.33 \pm 0.58/3$ | $2.67 \pm 0.58/3$ | $3.00 \pm 0.00/3$ |

The 384-candidate depth was the prespecified primary budget; 1 000, 2 000, and 5 000 were secondary hypothetical budgets. Deterministic controls are exact and have no stochastic variation. Learned-method values are mean ± sample standard deviation over seeds 42–44. Numerical ordering does not imply statistical significance.

#### Supplementary Table S6. Extended evaluation-panel clone rank audit

| Target | Clone | Total count | Count rank | Peak rank | Direct $O_i$ rank | Count–$O_i$ fusion | Tie-$O_i$ rank | Seed 42 | Seed 43 | Seed 44 |
|---|---|---|---|---|---|---|---|---|---|---|
| FAF1 | FAF1_AM_#8 | 7 | 10 704 | 16 213 | 16 252 | 14 705 | 11 680 | 1 501 | 1 199 | 1 746 |
| FAF1 | FAF1_AM_#20 | 891 | 3 | 3 | 3 | 3 | 3 | 257 | 233 | 136 |
| FAF1 | FAF1_AM_#10 | 38 | 588 | 390 | 466 | 502 | 590 | 5 658 | 4 845 | 6 145 |
| FAF1 | FAF1_AM_#12 | 6 | 14 939 | 11 156 | 12 459 | 14 977 | 14 836 | 3 837 | 3 801 | 4 298 |
| FAF1 | FAF1_#134 | 22 | 1 082 | 5 736 | 1 428 | 1 092 | 1 085 | 1 317 | 2 869 | 1 045 |
| FAF1 | FAF1_#138 | 6 | 12 677 | 8 306 | 10 626 | 12 476 | 13 471 | 361 | 317 | 359 |
| FAF1 | FAF1_#228 | 9 | 6 365 | 16 182 | 16 002 | 11 844 | 6 662 | 13 656 | 8 616 | 12 013 |
| VEGFR | AM-#3 | 6 638 | 6 | 10 | 10 | 8 | 6 | 4 | 2 186 | 1 244 |
| VEGFR | AM-#63 | 868 | 120 | 144 | 138 | 129 | 120 | 282 | 342 | 326 |
| VEGFR | AM-#83 | 371 | 330 | 312 | 312 | 318 | 329 | 87 | 197 | 224 |

Lower rank indicates higher prioritization. Count, peak, direct-$O_i$, count–$O_i$ fusion, and tie-$O_i$ ranks are deterministic; count–$O_i$ fusion is the unweighted sum of count rank and direct-$O_i$ rank, and tie-$O_i$ uses $O_i$ only within exact total-count ties. AbPACER columns report direct frozen ranks for the indicated seeds. Evaluation membership and apparent-affinity metadata are reported in Supplementary Table S3.

### 5 Representation, scoring/refinement, and frozen-output candidate-level analyses

All ablations use the fixed split, affinity-label-blind targets, seed-specific schedules, frozen candidate manifests, and direct model-score ranking. Ablated conditions modify the parent-aware representation or neural-scoring pathway while retaining the remaining common protocol.

#### Supplementary Table S7. Completed direct-score controlled ablations

Only completed, verified direct-score conditions are reported. All conditions used the fixed split, affinity-label-blind targets, seed-specific schedules, frozen candidate manifests, and direct model-score ranking.

*FAF1*

| Condition | Parameters | 384 | 2 000 | 5 000 |
|---|---|---|---|---|
| Full AbPACER | 1.378 M | 2.00 ± 0.00/7 | 3.67 ± 0.58/7 | 5.33 ± 0.58/7 |
| No query-level IgBERT context | 0.575 M | 1.67 ± 0.58/7 | 4.00 ± 1.00/7 | 5.67 ± 0.58/7 |
| Without local residual refinement | 1.361 M | 1.33 ± 0.58/7 | 3.67 ± 0.58/7 | 5.67 ± 0.58/7 |
| Without neighbor-derived forward features | 1.178 M | 1.00 ± 0.00/7 | 2.67 ± 1.53/7 | 5.00 ± 1.00/7 |

*VEGFR*

| Condition | Parameters | 384 | 2 000 | 5 000 |
|---|---|---|---|---|
| Full AbPACER | 1.378 M | 2.33 ± 0.58/3 | 2.67 ± 0.58/3 | 3.00 ± 0.00/3 |
| No query-level IgBERT context | 0.575 M | 2.67 ± 0.58/3 | 3.00 ± 0.00/3 | 3.00 ± 0.00/3 |
| Without local residual refinement | 1.361 M | 2.33 ± 0.58/3 | 3.00 ± 0.00/3 | 3.00 ± 0.00/3 |
| Without neighbor-derived forward features | 1.178 M | 0.67 ± 0.58/3 | 2.33 ± 0.58/3 | 3.00 ± 0.00/3 |

Parameters are rounded task-specific trainable counts and exclude frozen IgBERT. The no-query-level-IgBERT-context condition retained the IgBERT-derived neighbor graph and local-family supervision and was not a complete no-IgBERT experiment. The condition without local residual refinement used the neighbor-aware Base score directly. The condition without neighbor-derived forward features removed the learned neighbor representation and auxiliary neighborhood features from the forward path but retained local-family supervision and therefore was not fully neighbor-free.

Removing neighbor-derived forward features produced the clearest shared top-384 contribution across the two targets. Local residual refinement mainly affected FAF1 boundary-level recovery, whereas query-level IgBERT effects were target dependent. Because the ablations were not parameter matched, they support pathway-level interpretation rather than component-wise causal attribution. All 12 recomputed ablation runs that retained residual fitting (the no-query-level-IgBERT-context and without-neighbor-derived-forward-features conditions across two targets and three seeds) completed without nonfinite predictions or numerical collapse, and each selected a residual checkpoint that improved held-out anchor-pair loss relative to the zero-residual state. The no-residual condition did not fit or select a residual checkpoint by definition.

#### Supplementary Table S8. FAF1_#138 tie-aware pathway rank audit

The frozen-output audits reused the six seed-specific final AbPACER rankings and frozen top-5% manifests without new training, inference, checkpoint selection, calibration, score fusion, consensus ranking, or outcome-based tuning. Base-only and without-neighbor ranks came from their corresponding frozen stage and ablation outputs.

| Ranking rule or condition | Seed 42 | Seed 43 | Seed 44 | Top-384 inclusion |
|---|---|---|---|---|
| Deterministic total-count rank | 12 677 | 12 677 | 12 677 | 0/3 |
| Best possible rank under any within-count-6 tie-break | 11 698 | 11 698 | 11 698 | 0/3 |
| Without neighbor-derived forward features | 1 978 | 1 024 | 3 415 | 0/3 |
| Neighbor-aware Base score without residual refinement | 456 | 1 016 | 359 | 1/3 |
| Final AbPACER score | 361 | 317 | 359 | 3/3 |

FAF1_#138 had total count 6 and belonged to a 4 626-candidate tie spanning ranks 11 698–16 323. The count-6 tier began below the 384-candidate boundary, so no count-preserving tie ordering could place FAF1_#138 within the primary assay budget. The best-possible tie-break row is a mathematical bound rather than a fitted comparator. The Base score created the large

global promotion, and residual refinement stabilized top-384 inclusion across seeds. No other measured clone whose exact-count tie began outside the top 384 was promoted into the top 384 in any seed. The clone-level analysis was not used for model, checkpoint, or seed selection. Clone-level ranks for the complete evaluation panels are reported in Supplementary Table S6.

#### Supplementary Table S9. Measured-panel-excluded mutation-pattern audit

For target $t$ and seed $s$, let $A_s$ denote the final AbPACER top-384 set and $C$ the deterministic total-count top-384 set. AbPACER-only and count-only displaced candidates were $A_s \setminus C$ and $C \setminus A_s$, respectively. Evaluation-panel clone keys were removed before feature analysis. Each exact-substitution or position-level discovery feature was compared by a two-sided Fisher exact test, with Benjamini–Hochberg correction applied separately within each target × seed × feature-level family. The prespecified VH-position-57 F→L/I motif was a singleton family within each applicable target and seed. Stable features required enrichment in the AbPACER-only direction, BH FDR $\leq 0.05$, at least five AbPACER-only candidates carrying the feature within the corresponding target-by-seed comparison, and satisfaction of all three criteria in at least two seeds.

**Panel A. Predefined FAF1 VH-position-57 F→L/I motif**

| Seed | AbPACER-only selections | Count-only displaced candidates | Difference |
|---|---|---|---|
| 42 | 59.0% | 1.3% | +57.7 percentage points |
| 43 | 32.4% | 1.9% | +30.5 percentage points |
| 44 | 52.1% | 1.4% | +50.7 percentage points |

Evaluation-panel clones were excluded from both groups before feature analysis. The motif comprised parent-referenced F→L or F→I substitutions at VH position 57. After the mutation-analysis feature set was frozen, the motif was descriptively found in 5/7 FAF1 evaluation-panel clones, including FAF1_#138. This is a target-specific selection signature, not evidence of causality, and the comparison does not establish that unmeasured AbPACER-only candidates have superior affinity.

The AbPACER-only median count ranks for seeds 42, 43, and 44 were 4 884.5, 1 193.5, and 4 732.0, respectively, whereas the corresponding count-only displaced median ranks were 198.5, 212.5, and 193.0. This count-rank imbalance motivated a descriptive interpretation; the mutation comparison was not a count-independent mutation-effect test.

**Panel B. Stable feature-level discovery results**

| Target | Feature | Qualifying seeds | Mean AbPACER-only prevalence | Mean displaced prevalence | Mean difference | Minimum BH FDR |
|---|---|---|---|---|---|---|
| FAF1 | `VH:57:any_change` | 42,43,44 | 48.4% | 1.8% | +46.6 pp | $4.81 \times 10^{-61}$ |
| FAF1 | `VH:57:F>L` | 42,43,44 | 38.9% | 0.8% | +38.1 pp | $3.73 \times 10^{-46}$ |
| VEGFR | `VH:30:any_change` | 42,44 | 75.9% | 58.9% | +17.0 pp | $4.44 \times 10^{-25}$ |
| FAF1 | `VH:57:F>I` | 42,44 | 9.0% | 0.8% | +8.2 pp | $3.43 \times 10^{-8}$ |
| VEGFR | `VL:35:any_change` | 43,44 | 7.8% | 2.1% | +5.8 pp | $3.17 \times 10^{-3}$ |
| FAF1 | `VL:82:any_change` | 42,44 | 2.7% | 0.0% | +2.7 pp | $4.28 \times 10^{-2}$ |

The `VH:57:any_change`, `VH:57:F>L`, and `VH:57:F>I` rows are nested descriptions of the same mutation family and are not independent biological mechanisms. FAF1 `VL:82:any_change` is exploratory and cutoff-borderline, with seed-42 and seed-44 BH FDR values of 0.0428 and 0.0486 and AbPACER-only carrier counts of 9 and 9, respectively. None of the six signals establishes a causal affinity effect, cross-target generality, or superior apparent affinity among unmeasured AbPACER-only candidates.

*Audit note.* The frozen-input artifact gate passed 53/53 checks, and measured-panel apparent-affinity data were loaded only after the feature set had been frozen. Complete candidate-level mutation-token results are retained as machine-readable artifacts in the versioned archival release.

### 6 Comparator adaptations and task-specific parameter accounting

Parameter counts report unique task-specific trainable parameters updated at any stage and exclude frozen feature extractors where stated. Comparator results are common-protocol adaptations or reimplementations rather than exact reproductions of original tasks. The original Ens-Grad task varied CDR-H3 within an otherwise fixed antibody framework, whereas the present comparator ranked paired scFv candidates carrying parent-relative changes across VH and VL. AbAffinity was adapted from quantitative affinity prediction to the common $O_i$-based pairwise ranking objective. A2Binder-HL branch use and antigen-length constraints are detailed in Panel B.

#### Supplementary Table S10. Comparator adaptations and task-specific parameter accounting

**Panel A. Phage-display task-specific parameter summary**

| Method | Target | Task-specific trainable parameters |
|---|---|---|
| Ens-Grad CNN | FAF1 | 58 062 |
| Ens-Grad CNN | VEGFR | 58 062 |
| A2Binder-HL | FAF1 | 114 631 425 |
| A2Binder-HL | VEGFR | 114 631 425 |
| AbAffinity | FAF1 | 651 044 535 |
| AbAffinity | VEGFR | 651 044 535 |
| AbPACER | FAF1 | 1 378 348 |
| AbPACER | VEGFR | 1 378 396 |

Trainable counts for AbPACER sum the unique base and bounded-correction parameters updated at different stages and exclude frozen IgBERT.

**Panel B. A2Binder branch usage**

| Benchmark | A2Binder branches used | Interpretation |
|---|---|---|
| Phage-display | Released heavy- and light-chain branches; antigen branch not used | A2Binder-HL antibody-backbone adaptation. The released antigen pathway accepted 300 tokens, corresponding to at most 298 antigen residues after special tokens. It would therefore truncate 352 of 650 FAF-1 residues and 447 of 745 VEGFR-2 residues. Full-length input would require a nonreleased modification of the antigen CNN input shape or feature mapping; this analysis is not a full antigen-aware A2Binder reproduction. |
| AlphaSeq | Released heavy-, light-, and antigen branches with fixed peptide `PDVDLGDISGINAS` | Antigen-aware common-split adaptation to the public HR2 landscape. |

The antigen lengths refer to the experimental protein regions after exclusion of the FAF-1 construct-derived linker/Myc/His tail and the VEGFR-2 Fc fusion. No truncated-antigen or custom length-expanded phage model was trained; the reported A2Binder-HL parameter count, predictions, rankings, and recovery values are unchanged.

The AlphaSeq common-split training protocol, target transformation, fit-only neighbor-reference policy, bandwidth selection, and test-label firewall are detailed in Section 1.6.

**Panel C. AlphaSeq task-specific parameter summary**

| Model | Task-specific trainable parameters |
|---|---|
| AbPACER-MSE | 1 378 348 |
| AbAffinity | 651 044 535 |
| A2Binder | 263 850 074 |
| Ens-Grad CNN | 58 062 |

AlphaSeq parameter counts are per-seed totals across all deployed models or ensemble constituents and include every unique task-specific parameter updated at any training stage; frozen or non-trainable parameters are excluded. The pooled AbPACER-MSE value of 1 378 348 counts every unique parameter updated during its base or residual stage and excludes frozen IgBERT; the residual-stage snapshot contains 17 289 currently trainable parameters and 1 361 059 base parameters frozen at that stage. The AbAffinity and A2Binder values are pooled single-model totals; the A2Binder value excludes 65 536 non-trainable parameters. The Ens-Grad CNN value of 58 062 sums all six pooled ensemble constituents.

Validation-based training sufficiency of the primary AbPACER runs and principal high-capacity comparators is summarized in Supplementary Table S15.

### 7 Detailed AlphaSeq results and sensitivity analyses

All analyses followed the immutable fit/validation/test split and test-label firewall in Section 1.6, used seeds 42–44 and frozen predictions, and involved no new training, checkpoint reselection, or recalibration. Primary analyses used pooled-model predictions; the training-granularity sensitivity additionally reused frozen predictions from the four library-specific AbPACER-MSE models. AbPACER-MSE used the appropriate library-specific parental sequence and within-library feature and neighbor providers. The 384-selection depth matched the primary fixed-budget benchmark used in the phage-display arm, although supervision and recovery denominators remained distinct.

#### 7.1 Seed-specific global and fixed-budget results

The primary fixed-budget endpoint was overlap between each predicted top 384 and the measured true top 384. Top-1%, top-5%, and top-10% recovery among the same predictions remained secondary threshold-based analyses.

#### Supplementary Table S11. Seed-specific AlphaSeq results

**Panel A. Global regression metrics**

| Model | Seed | Pearson | Spearman | RMSE | MAE |
|---|---|---|---|---|---|
| AbPACER-MSE | 42 | 0.690695 | 0.654166 | 0.967853 | 0.737690 |
| AbPACER-MSE | 43 | 0.685127 | 0.650961 | 0.975535 | 0.743060 |
| AbPACER-MSE | 44 | 0.686416 | 0.649897 | 0.974859 | 0.741394 |
| AbAffinity | 42 | 0.687319 | 0.653636 | 0.977897 | 0.741334 |
| AbAffinity | 43 | 0.683069 | 0.651707 | 0.975986 | 0.745737 |
| AbAffinity | 44 | 0.686594 | 0.651013 | 0.971542 | 0.743199 |
| A2Binder | 42 | 0.678025 | 0.647686 | 1.025987 | 0.769571 |
| A2Binder | 43 | 0.676887 | 0.644881 | 0.987545 | 0.748704 |
| A2Binder | 44 | 0.672563 | 0.642203 | 1.003416 | 0.757942 |
| Ens-Grad CNN | 42 | 0.630684 | 0.601092 | 1.090084 | 0.880884 |
| Ens-Grad CNN | 43 | 0.635763 | 0.606602 | 1.086437 | 0.876926 |
| Ens-Grad CNN | 44 | 0.634473 | 0.605080 | 1.086391 | 0.878616 |

**Panel B. Primary top-384 overlap and secondary threshold-based recovery among 384 selected variants**

| Model | Seed | True top-384 hits | Top-1% hits | Top-5% hits | Top-10% hits |
|---|---|---|---|---|---|
| AbPACER-MSE | 42 | 188 | 88 | 239 | 306 |
| AbPACER-MSE | 43 | 189 | 89 | 237 | 302 |
| AbPACER-MSE | 44 | 184 | 89 | 236 | 299 |
| AbAffinity | 42 | 190 | 91 | 238 | 304 |
| AbAffinity | 43 | 189 | 83 | 239 | 303 |
| AbAffinity | 44 | 185 | 85 | 242 | 304 |
| A2Binder | 42 | 173 | 80 | 225 | 289 |
| A2Binder | 43 | 183 | 79 | 229 | 292 |
| A2Binder | 44 | 171 | 79 | 226 | 291 |
| Ens-Grad CNN | 42 | 158 | 71 | 201 | 274 |
| Ens-Grad CNN | 43 | 147 | 65 | 198 | 280 |
| Ens-Grad CNN | 44 | 157 | 70 | 204 | 278 |

True top-384 overlap is the primary fixed-budget endpoint. The top-1%, top-5%, and top-10% values are secondary threshold-based analyses. Every value counts members recovered among the same 384 selected predictions. Exact ties were resolved using the deterministic fixed-test row key.

#### 7.2 Library-stratified regression

The frozen pooled predictions were evaluated separately for 14H, 14L, 91H, and 95L.

#### Supplementary Table S12. AlphaSeq library-stratified three-seed regression results

Values are mean ± sample standard deviation over fixed seeds 42, 43, and 44. Each library used identical fixed-test membership across methods and seeds.

| Model | Library | Test variants | Pearson | Spearman | RMSE | MAE |
|---|---|---|---|---|---|---|
| AbPACER-MSE | 14H | 2 290 | 0.636 ± 0.003 | 0.524 ± 0.003 | 0.999 ± 0.004 | 0.789 ± 0.005 |
| AbPACER-MSE | 14L | 3 048 | 0.645 ± 0.005 | 0.650 ± 0.005 | 1.159 ± 0.007 | 0.886 ± 0.005 |
| AbPACER-MSE | 91H | 2 162 | 0.525 ± 0.007 | 0.408 ± 0.006 | 1.075 ± 0.006 | 0.826 ± 0.005 |
| AbPACER-MSE | 95L | 4 170 | 0.658 ± 0.002 | 0.614 ± 0.002 | 0.721 ± 0.002 | 0.564 ± 0.002 |
| AbAffinity | 14H | 2 290 | 0.637 ± 0.007 | 0.531 ± 0.003 | 1.003 ± 0.009 | 0.798 ± 0.008 |
| AbAffinity | 14L | 3 048 | 0.651 ± 0.003 | 0.656 ± 0.005 | 1.156 ± 0.009 | 0.882 ± 0.003 |
| AbAffinity | 91H | 2 162 | 0.503 ± 0.005 | 0.398 ± 0.003 | 1.088 ± 0.005 | 0.842 ± 0.003 |
| AbAffinity | 95L | 4 170 | 0.660 ± 0.001 | 0.620 ± 0.001 | 0.720 ± 0.004 | 0.561 ± 0.003 |
| A2Binder | 14H | 2 290 | 0.621 ± 0.004 | 0.523 ± 0.002 | 1.044 ± 0.016 | 0.818 ± 0.011 |
| A2Binder | 14L | 3 048 | 0.628 ± 0.001 | 0.631 ± 0.001 | 1.204 ± 0.024 | 0.906 ± 0.008 |
| A2Binder | 91H | 2 162 | 0.513 ± 0.003 | 0.411 ± 0.002 | 1.091 ± 0.014 | 0.836 ± 0.009 |
| A2Binder | 95L | 4 170 | 0.656 ± 0.001 | 0.615 ± 0.001 | 0.744 ± 0.027 | 0.579 ± 0.017 |
| Ens-Grad CNN | 14H | 2 290 | 0.547 ± 0.006 | 0.473 ± 0.001 | 1.146 ± 0.007 | 0.940 ± 0.009 |
| Ens-Grad CNN | 14L | 3 048 | 0.602 ± 0.003 | 0.610 ± 0.002 | 1.268 ± 0.007 | 1.061 ± 0.008 |
| Ens-Grad CNN | 91H | 2 162 | 0.416 ± 0.006 | 0.368 ± 0.008 | 1.226 ± 0.006 | 1.016 ± 0.008 |
| Ens-Grad CNN | 95L | 4 170 | 0.611 ± 0.002 | 0.573 ± 0.001 | 0.796 ± 0.004 | 0.641 ± 0.002 |

#### 7.3 Within-library ranking and training-granularity sensitivity

Within-library comparisons reduced the influence of library size, difficulty, and cross-library prediction scale. The 10×-confident ROC–AUC retained pairs separated by at least one $\log_{10}(K_D)$ unit in both orientations; all-pair concordance gave half credit to predicted ties, and Kendall's $\tau_b$ accounted for ties. Metrics were calculated by seed within each library and macro-averaged across 14H, 14L, 91H, and 95L. This AbRank-inspired evaluation used neither the AbRank dataset nor its splits.

Aggregate pooled-model metrics are reported in Main Table 4; the supplementary analyses below provide the seed-specific, library-stratified, and training-granularity results not shown in the main manuscript.

#### Supplementary Table S13. Global and within-library training-granularity sensitivity

##### Panel A. Global and secondary threshold-based training-granularity sensitivity

| Configuration | Parameters | Pearson | Spearman | RMSE | MAE | Top-1% | Top-5% | Top-10% |
|---|---|---|---|---|---|---|---|---|
| Pooled shared model | 1.378 M | 0.687 ± 0.003 | 0.652 ± 0.002 | 0.973 ± 0.004 | 0.741 ± 0.003 | 88.7 ± 0.6 | 237.3 ± 1.5 | 302.3 ± 3.5 |
| Four library-specific models | 5.513 M | 0.687 ± 0.001 | 0.651 ± 0.001 | 0.973 ± 0.001 | 0.741 ± 0.001 | 86.3 ± 0.6 | 230.3 ± 6.4 | 297.7 ± 4.0 |

Values are mean ± sample standard deviation over seeds 42–44. The pooled and library-specific configurations had essentially identical global regression performance. The pooled model used one quarter of the task-specific parameters and showed numerically higher secondary threshold-based recovery across the reported top-1%, top-5%, and top-10% sets. These numerical differences were not interpreted as statistical separation.

##### Panel B. Within-library pairwise training-granularity sensitivity

| Configuration | 10× AUC | Concordance | Kendall $\tau_b$ |
|---|---|---|---|
| Pooled shared model | 0.877 ± 0.002 | 0.694 ± 0.001 | 0.388 ± 0.002 |
| Four library-specific models | 0.877 ± 0.001 | 0.694 ± 0.001 | 0.387 ± 0.001 |

Panel B values are macro-averages across 14H, 14L, 91H, and 95L, reported as mean ± sample standard deviation over seeds 42–44. Pairwise ranking metrics were essentially unchanged by training granularity. These numerical differences were not interpreted as statistical separation.

### 8 Manifest provenance, cohort definition, and release scope

The AlphaSeq benchmark used the 71,834-variant cohort processed for AbAffinity, with identical immutable memberships across methods. The phage analyses used frozen target-specific candidate manifests.

#### Supplementary Table S14. Cohort and frozen-manifest provenance summary

| Resource | Rows/candidates | Evaluation role | Frozen membership |
|---|---|---|---|
| AlphaSeq fit | 51 139 | Supervised fitting | Fixed |
| AlphaSeq validation | 9 025 | Bandwidth and checkpoint selection | Fixed |
| AlphaSeq test | 11 670 | Final evaluation | Fixed |
| FAF1 top-5% manifest | 16 323 | Primary phage candidate universe | Yes |
| VEGFR top-5% manifest | 7 487 | Primary phage candidate universe | Yes |
| Top-1%/top-10% manifests | See Table S2 | Reference manifest definitions | Yes |

The versioned archival release contains the frozen manifests, complete seed-specific predictions and rankings, resolved configurations, software-environment records, and file-level SHA-256 checksums.

The planned archival release will be linked through the AbPACER project repository and its Zenodo archive described in the main manuscript's Availability of data and materials section. The original AlphaSeq sequence and measured-label cohort will not be redistributed; AlphaSeq-derived release items will be limited to label-stripped predictions, derived evaluation outputs, row identifiers or hashes required for provenance, and links to the upstream AlphaSeq Zenodo record cited in the main manuscript. Software will be released under the BSD-3-Clause license, and phage-display data and paper-associated artifacts under CC BY 4.0. Raw PacBio reads are associated with NCBI BioProject PRJNA1506661 and will be released through the NCBI Sequence Read Archive no later than publication, while processed phage data and reproducibility artifacts will be archived in Zenodo. Individual SRA run accessions and the AbPACER Zenodo DOI will be added when assigned.

### 9 Validation-based training-sufficiency and checkpoint audit

The audit covered the primary AbPACER phage runs, the pooled AbPACER-MSE AlphaSeq runs, and the principal high-capacity comparators. It tested whether the reported checkpoints were validation selected, observed for sufficient additional epochs, and free of numerical collapse. No retrospective recovery outcome was used for checkpoint reselection. Complete seed-, ensemble-member-, and epoch-level checkpoint histories are provided in the versioned archival release; the table below summarizes the validation-selected training durations used in the manuscript analyses.

#### Supplementary Table S15. Validation-based training sufficiency

**Panel A. AbPACER phage direct-ranking runs**

| Target | Seed | Base maximum | Base executed | Selected Base | Best validation Spearman | Residual maximum | Residual executed | Selected residual | Best residual validation loss | Improvement over epoch 0 | Checkpoint evidence | Collapse |
|---|---|---|---|---|---|---|---|---|---|---|---|---|
| FAF1 | 42 | 20 | 13 | 10 | 0.1628 | 15 | 11 | 8 | 0.6989 | 0.0770 | Validated improvement | No |
| FAF1 | 43 | 20 | 11 | 8 | 0.1842 | 15 | 11 | 8 | 0.7264 | 0.0679 | Validated improvement | No |
| FAF1 | 44 | 20 | 15 | 12 | 0.1822 | 15 | 11 | 8 | 0.6997 | 0.0635 | Validated improvement | No |
| VEGFR | 42 | 20 | 10 | 7 | 0.4249 | 15 | 9 | 6 | 0.7452 | 0.1363 | Validated improvement | No |
| VEGFR | 43 | 20 | 4 | 1 | 0.3993 | 15 | 9 | 6 | 0.7297 | 0.1234 | Validated improvement | No |
| VEGFR | 44 | 20 | 4 | 1 | 0.3798 | 15 | 9 | 6 | 0.7596 | 0.1721 | Validated improvement | No |

Base checkpoints used validation Spearman correlation with $O_i$, and residual checkpoints used held-out local anchor-pair loss. Displayed validation values are rounded to four decimal places; exact values and source paths are retained in the audit trace. Improvement over epoch 0 is the absolute reduction in held-out anchor-pair loss. Residual epoch 0 was eligible; every run selected a nonzero checkpoint with validated improvement. Each Base and residual holdout fit continued for three post-best epochs and passed its search-window and numerical-health checks. Base holdout training and final refitting used FP32.

**Panel B. Comparator checkpoint sufficiency summarized across seeds 42–44**

| Model | Benchmark/target | Maximum scheduled epoch | Selected epoch range | Post-best epoch range | Validation criterion | Collapse | Stopping evidence |
|---|---|---|---|---|---|---|---|
| A2Binder-HL | FAF1 | 20 | 6–11 | 3 | Validation Spearman with $O_i$ | No | Patience exhausted |
| A2Binder-HL | VEGFR | 20 | 5–9 | 3 | Validation Spearman with $O_i$ | No | Patience exhausted |
| AbAffinity | FAF1 | 20 | 9–10 | 3 | Validation Spearman with $O_i$ | No | Patience exhausted |
| AbAffinity | VEGFR | 20 | 6–8 | 3 | Validation Spearman with $O_i$ | No | Patience exhausted |
| A2Binder | Pooled AlphaSeq | 12 | 2 | 10 | Global validation Pearson | No | Fixed horizon completed |
| AbAffinity | Pooled AlphaSeq | 100 | 10–11 | 89–90 | Global validation Pearson | No | Fixed horizon completed |

Ranges are the observed minima and maxima over the three fixed seeds. Post-best epochs equal executed minus selected epoch. Phage comparator early stopping used patience three; AlphaSeq comparators completed their fixed horizons. No principal comparator showed prediction collapse or evidence of training-horizon truncation.

**Panel C. Pooled AbPACER-MSE AlphaSeq runs**

| Seed | Base maximum / executed epoch | Selected Base epoch | Base post-best epochs | Residual maximum / executed epoch | Selected residual epoch | Residual post-best epochs |
|---|---|---|---|---|---|---|
| 42 | 100 / 25 | 15 | 10 | 100 / 15 | 5 | 10 |
| 43 | 100 / 26 | 16 | 10 | 100 / 13 | 3 | 10 |
| 44 | 100 / 27 | 17 | 10 | 100 / 16 | 6 | 10 |

Pooled AbPACER-MSE used one shared model per seed across the 14H, 14L, 91H, and 95L libraries. Both Base and residual checkpoints were selected by maximizing a single global Pearson correlation over the 9 025 pooled validation rows, with patience-10 early stopping. Base parameters remained frozen during residual fitting, and test labels were not used for checkpoint selection. The maximum/executed entries distinguish the configured maximum horizon from the epochs actually completed before early stopping. No separate post-selection refitting was performed; the frozen manuscript predictions were linked to the validation-selected checkpoints through the freeze and final-evaluation manifests. All three seeds had verified checkpoint and blind-prediction provenance, with no selected-epoch conflict.

The Ens-Grad-inspired CNN was retained as a lightweight architectural reference. Its frozen equal-weight ensemble outputs were evaluated without member removal, checkpoint replacement, or outcome-based reselection.

This audit addresses validation-selected training sufficiency and numerical health rather than statistical significance.
